# Visuomotor anomalies in achiasmatic mice expressing a transfer-defective Vax1 mutant

**DOI:** 10.1101/2020.10.20.346551

**Authors:** Kwang Wook Min, Namsuk Kim, Jae Hoon Lee, Younghoon Sung, Museong Kim, Eun Jung Lee, Jong-Myeong Kim, Jae-Hyun Kim, Jaeyoung Lee, Wonjin Cho, Jee Myung Yang, Nury Kim, Jaehoon Kim, C. Justin Lee, Young-Gyun Park, Seung-Hee Lee, Han-Woong Lee, Jin Woo Kim

## Abstract

In binocular animals that exhibit stereoscopic visual responses, the axons of retinal ganglion cells (RGCs) connect to brain areas bilaterally by forming a commissure called the optic chiasm (OC). Ventral anterior homeobox 1 (Vax1) contributes to formation of the OC, acting endogenously in optic pathway cells and exogenously in growing RGC axons. Here, we generated *Vax1^AA/AA^* mice expressing the Vax1^AA^ mutant, which is incapable of intercellular transfer. We found that RGC axons cannot take up Vax1^AA^ protein from *Vax1^AA/AA^* mouse optic stalk (OS) cells, of which maturation is delayed, and fail to access the midline. Consequently, RGC axons of *Vax1^AA/AA^* mice connect exclusively to ipsilateral brain areas, resulting in the reduced visual acuity and the abnormal oculomotor responses. Together, our study provides physiological evidences for the necessity of intercellular transfer of Vax1 and the importance of the bilateral RGC axon projection in visuomotor responses.

## Introduction

Animals collect visual information by the eyes, where photoreceptors in the retina convert light stimuli into electrochemical signals (Seabrook et al., 2017). The signals are then transmitted to inner retinal neural circuits before being sent to the brain via retinal ganglion cells (RGCs). RGC axons, which are bundled in the optic nerve, deliver visual information to multiple brain areas, including the dorsal lateral geniculate nucleus (dLGN) of the thalamus, for pattern and color recognition; the superior colliculus (SC) of the midbrain, for oculomotor responses; and the suprachiasmatic nucleus (SCN) of the hypothalamus, for circadian rhythm control (Rusak and Groos, 1982; Seabrook et al., 2017; Zhang et al., 2017).

In many binocular animals, RGC axons are not only wired to brain areas on the same (i.e., ipsilateral) side but are also connected to those on the opposite (i.e., contralateral) side (Herrera et al., 2019; Petros et al., 2008). The population of RGCs whose axons project to the ipsilateral brain areas is variable among vertebrate species. Human RGC axons are split equally to both sides at the midline, whereas all RGC axons extend exclusively to the contralateral side across the midline in *Xenopus laevis* and zebrafish (Herrera et al., 2019; Petros et al., 2008). In mice, only a minority of RGCs connects to the ipsilateral brain areas: ∼3% in pigmented mice and ∼1% in albino mice (Rice et al., 1995).

RGC axons form a midline structure called the optic chiasm (OC), which is located beneath the SCN and splits the axon bundles into ipsilateral and contralateral paths (Herrera et al., 2019; Petros et al., 2008). Pathway selection for RGC axons at the OC is determined by specific guidance cues. For example, Ephrin-B2 and -B3 expressed in radial glia of the ventral hypothalamus (vHT) act through a receptor, EphB1, in RGC axons from the ventral and temporal (VT) retina to repel the axons from the midline, guiding their growth ipsilaterally (Williams et al., 2003). EphB1 is present in about 50% of human RGCs and ∼3% of mouse RGCs, but is absent in *Xenopus* and zebrafish RGCs, suggesting a critical role of EphB1 in ipsilateral pathway selection by RGC axons (Herrera et al., 2003; Rebsam et al., 2012). Pathway-selection cues are not only provided by cells located along RGC axon growth tracks, but also by neighboring RGC axons. Sonic hedgehog (Shh), which is expressed in contralaterally projecting RGC axons, also serves as a repulsive cue in the OC by acting on its co-receptor, Boc, which is expressed in RGC axons from the VT mouse retina (Peng et al., 2018).

The cues that guide the majority of mouse RGC axons across the midline, however, have not been identified as clearly as the ipsilateral guidance cues. Binding of vascular endothelial growth factor-a (Vegfa) to its receptor, neuropilin-1 (Nrp1), has been demonstrated to support the growth of RGC axons at the midline (Erskine et al., 2011). Homophilic interactions between neuronal cell adhesion molecule (Nr-CAM) expressed in RGC axons and vHT cells have also been suggested to promote midline crossings of RGC axons (Williams et al., 2006). Nr-CAM also cooperates with plexinA1, a receptor for semaphorin 6D (Sema6D), to support contralateral RGC axon projection (Kuwajima et al., 2012). However, a majority of RGC axons still cross the midline in mice lacking these cues, suggesting the presence of other key regulator(s) of contralateral RGC axon growth.

Ventral anterior homeobox 1 (Vax1) is expressed in ventral and medial regions of the vertebrate forebrain (Bertuzzi et al., 1999; Hallonet et al., 1999). The forebrain commissural structures, including the anterior commissure (AC), corpus callosum (CC), hippocampal commissure (HC), and OC, do not form properly in humans and mice having homozygous *VAX1* mutations (Bertuzzi et al., 1999; Mui et al., 2005; Slavotinek et al., 2012). Given the absence of *VAX1* gene expression in the commissural neurons, it had been thought that VAX1 functions as a transcription factor that induces the expression of axon growth factors in cells located along commissural axon growth tracks. However, it was found that the transcription factor activity of Vax1 is dispensable for the promotion of mouse RGC axon growth (Kim et al., 2014). More surprisingly, Vax1 protein was detected in mouse RGC axons, despite the absence of autonomous *Vax1* gene expression in RGCs (Kim et al., 2014). It was further found that Vax1 protein in mouse RGC axons is transferred from cells along RGC axon growth tracks and promotes the axon growth by stimulating local mRNA translation (Kim et al., 2014). However, whether intercellular Vax1 transfer is also critical for the growth of RGC axons *in vivo* has been remained unanswered.

Thus, we generated *Vax1^AA/AA^* mice, in which *Vax1* was replaced by a transfer-defective *Vax1^AA^* mutant. We found that Vax1^AA^ protein was incapable of binding to heparan sulfate proteoglycans (HSPGs) and penetrating RGC axons, resulting in retarded growth of RGC axons. Most of all, the OC was not formed in *Vax1^AA/AA^* mutant mice, and RGC axons projected exclusively to ipsilateral brain areas. Consequently, the *Vax1^AA/AA^* mice exhibited abnormal visuomotor responses, suggesting the importance of bilateral RGC axon growth in mice. Our findings in *Vax1^AA/AA^* mice, therefore, not only confirm the physiological importance of Vax1 transfer but also provide a molecular basis for the visuomotor anomalies seen in achiasmatic mammals.

## Results

### Identification of a CAG sugar-binding motif in Vax1

Vax1 was found to be secreted from cells that interact with developing mouse RGC axons and enters into the axons. Binding of Vax1 to RGC axons was mediated by heparan sulfate (HS) sugar chains of HSPGs, such as syndecan (Sdc) and glypican (Glp) (Kim et al., 2014). Another secreted homeodomain protein, Otx2 (orthodenticle homeobox 2), was also found to bind chondroitin sulfate (CS) sugar chains of chondroitin sulfate proteoglycan (Beurdeley et al., 2012; Miyata et al., 2012). It was further identified that binding of Otx2 to CS was mediated by the conserved glycosaminoglycan (GAG) binding motifs, [-X-B-B-X-B-X-] and [-X-B-B-B-X-X-B-X-] (Cardin and Weintraub, 1989).

We identified mouse Vax1 also contains a GAG-binding motif located at amino acids 100–105 (Figure 1A). To investigate the possibility that Vax1 binds to HS chains through this putative CAG-binding motif, we replaced lysine and arginine (KR) amino acid residues in the motif to two alanines (A), yielding Vax1^AA^ (Figure 1A). HeLa cells were transfected with DNA construct that expresses an mRNA encoding *Vax1* or *Vax1^AA^* together with enhanced green fluorescent protein (EGFP) but produces EGFP separately from Vax1 or Vax1^AA^ using an internal ribosome entry site (IRES). The KR-to-AA mutation suppressed Vax1 transfer from EGFP-positive donor cells to EGFP-negative recipient cells (Figure 1B). However, it did not significantly affect Vax1 transcription factor activity, which was assessed by monitoring the expression of a luciferase reporter downstream of a Vax1 target sequence in *transcription factor 7-like 2* (*Tcf7l2*) gene (Vacik et al., 2011) (Figure 1C). This finding contrasts with the reduced transcription factor activity of the Vax1^R152S^ mutant (Figure 1C), in which arginine 152 (R152) of the DNA binding motif was changed to serine (S) (Slavotinek et al., 2012).

**Figure 1.**
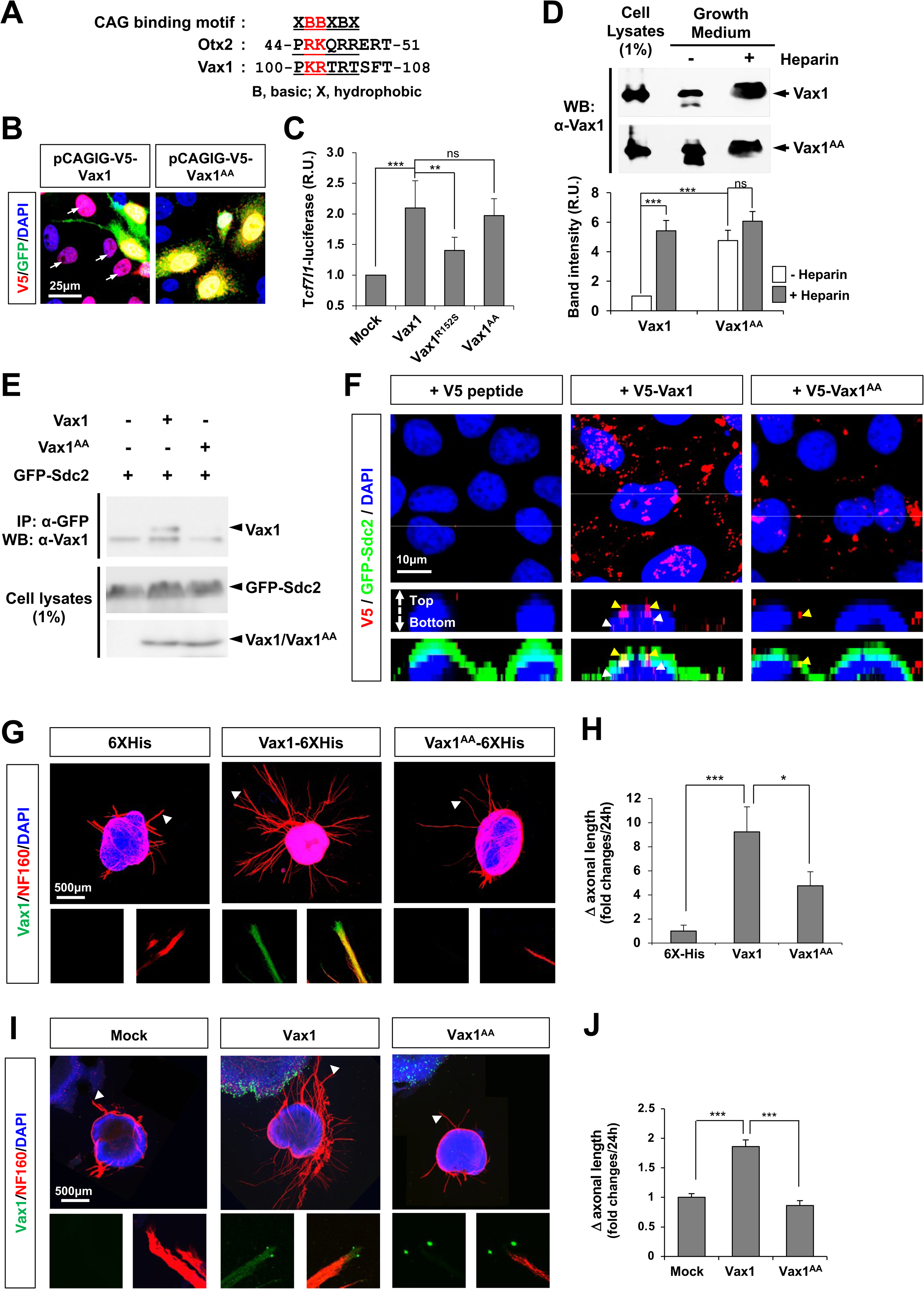
Identification a CAG sugar binding motif of Vax1. (**A**) Consensus amino acid sequences of GAG sugar-binding motifs in mouse Otx2 and Vax1. (**B**) Expression of V5- Vax1 and EGFP, which are independently translated from a same transcript, was examined by immunostaining of transfected HeLa cells with mouse anti-V5 (red) and chicken anti-GFP (green) antibodies. Arrows point HeLa cells express V5-Vax1 without EGFP, implicating the transfer of V5-Vax1 but not EGFP from V5;EGFP double-positive cells. (**C**) HEK293T cells were transfected with a DNA vector encodes Vax1, Vax1(R152S), or Vax1(KR/AA) cDNA together with a Vax1 target *Tcf7l2*-luciferase reporter DNA construct. Luciferase activities in the transfected cells were measured at 24- h post-transfection. The values are average obtained from four independent experiments and error bars denote standard deviations (SD). *p*-values were determined by ANOVA test (*, p<0.01; **, p<0.005; ***, p<0.001; ns, not significantly different). (**D**) V5-tagged Vax1 or Vax1^AA^ proteins were expressed in HEK293T cells, and growth media of the transfected cells were then collected after incubating for 3h in the presence (+) or absence (-) of heparin (10 mg/ml final; see Methods for details). Cell lysates and TCA-precipitated fractions of growth medium were analyzed by 10% SDS-PAGE and subsequent western blotting (WB) with anti-Vax1 antibody (α-Vax1). A graph below the WB data shows the relative intensities of Vax1 bands in the blots. The values are average obtained from four independent experiments and error bars denote SD. (**E**) Interactions of V5-Vax1 and V5- Vax1^AA^ with GFP-Sdc2 in HEK293T cells were assessed by immunoprecipitation (IP) with α-V5 and subsequent WB with α-GFP. Relative amounts of V5-Vax1 and GFP-Sdc2 in the cell lysates were also examined by WB. (**F**) V5-Vax1 or V5-Vax1^AA^ recombinant proteins were added into growth medium of HeLa cells expressing GFP-Sdc2 and incubated for 3 h. Vax1 proteins inside cells and/or at the cell surface were detected by immunostaining with mouse α-V5 (red) and chick α-GFP (green). **(G)** Retinas were isolated from E13.5 mice and cultured as described in Methods. Axonal lengths of retinal explants were measured at 24 h post-culture; then, the explants were treated with 6X-His-tagged recombinant Vax1 or Vax1^AA^ proteins. **(I)** Alternatively, retinal explants were co-cultured with HEK293T cells transfected with pCAGIG (Mock), pCAGIG-V5-Vax1, or pCAGIG-V5- Vax1^AA^. Axonal lengths were re-measured after 24 h and the explants were immunostained with α-Vax1 (green) and α-NF160 (red). Arrowheads indicate the areas magnified in each inset. (**H** and **J**) The changes in axonal length during the 24-h incubation period were shown in graphs. The values in the graph are averages and error bars denote SDs (n=6).

We found that the level of Vax1^AA^ protein in the growth media of the transfected cells was significantly elevated compared with that of Vax1 (Figure 1D). However, the amount of Vax1^AA^ in the medium was not further increased by the addition of free heparin, which competes with HS chains of HSPGs to bind extracellular Vax1 and release it from the HSPG-enriched cell surface into the growth medium (Lee et al., 2019) (Figure 1D). The results suggest reduced affinity of Vax1^AA^ for HS sugars compared with Vax1, together with a result that shows the impaired interaction between Vax1^AA^ and syndecan-2 (Sdc2) HSPG (Figure 1E). Consequently, Vax1^AA^ added to the growth medium could bind or penetrate HeLa cells much less efficiently than Vax1 (Figure 1F).

We, next, tested whether the KR-to-AA mutation also influences the binding and penetration of Vax1 to RGC axons and subsequent axon growth stimulation by Vax1. We found that Vax1^AA^ added into the growth medium of mouse retinal explants was not detected in neurofilament 160 (Nf160)-positive RGC axons nor did it induce axonal growth as efficiently as Vax1, which penetrated retinal axons and significantly promoted axonal growth (Figure 1G and 1H). We also found that Vax1^AA^ was not transferred from human embryonic kidney 293T cells to RGC axons, which were projected from co-cultured with mouse retinal explants (Figure 1I and 1J). Consequently, the lengths of RGC axons extending toward Vax1^AA^-expressing 293T cells were shorter than those growing towards Vax1-expressing 293T cells. These results suggest that the KR residues are necessary for the binding of Vax1 to HSPGs and subsequent penetration of Vax1 into RGC axons.

### Generation of Vax1^AA/AA^ mice

To investigate the consequences of the KR to AA mutation *in vivo*, we next introduced the corresponding mutation into the mouse *Vax1* gene to generate *Vax1^AA^* mice (Figure 1 - figure supplement 1A and B; see Methods for details). *Vax1^AA/AA^* homozygous mice were born without any recognizable morphological defects and survived without significant health problems, whereas the mice carry homozygous non-sense mutations (*Vax1^-/-^*) or hemizygous KR-to-AA mutation together non-sense mutation (*Vax1^AA/-^*) die after birth with the cleft palates that interfere with breathing of the mice (Bertuzzi et al., 1999) (Figure 1 - figure supplement 1C – E). Noticeably, the incisors grow continuously in approximately a quarter of *Vax1^AA/AA^* mice (Figure 1 - figure supplement 1D). The outgrowing incisors make the mice difficult to consume chow and cause a lethality after weaning, unless the outgrown incisors were not cut off regularly (Figure 1 - figure supplement 1E).

Using *Vax1^AA/AA^* mouse embryos, we examined whether the mutation affects intercellular transfer of Vax1 *in vivo*. We found Vax1^AA^ was strongly expressed in ventral medial forebrain structures, including the optic stalk (OS), of *Vax1^AA/AA^* mouse embryos at 14.5 days post-coitum (dpc; E14.5), a distribution pattern similar to that of Vax1 in *Vax1^+/+^* littermate mice (Figure 2A). Unlike Vax1, which was present in RGC axons as well as OS cells in *Vax1^+/+^* mice, Vax1^AA^ was not detectable in the RGC axons in *Vax1^AA/AA^* mice (Figure 2A). However, *Vax1^AA^* mRNA was present in the OS cells of *Vax1^AA/AA^* mice (Figure 2B), suggesting that the KR-to-AA mutation did not affect *Vax1* gene expression in the OS cells but interfered with Vax1 protein transfer from the OS cells to RGC axons.

**Figure 2.**
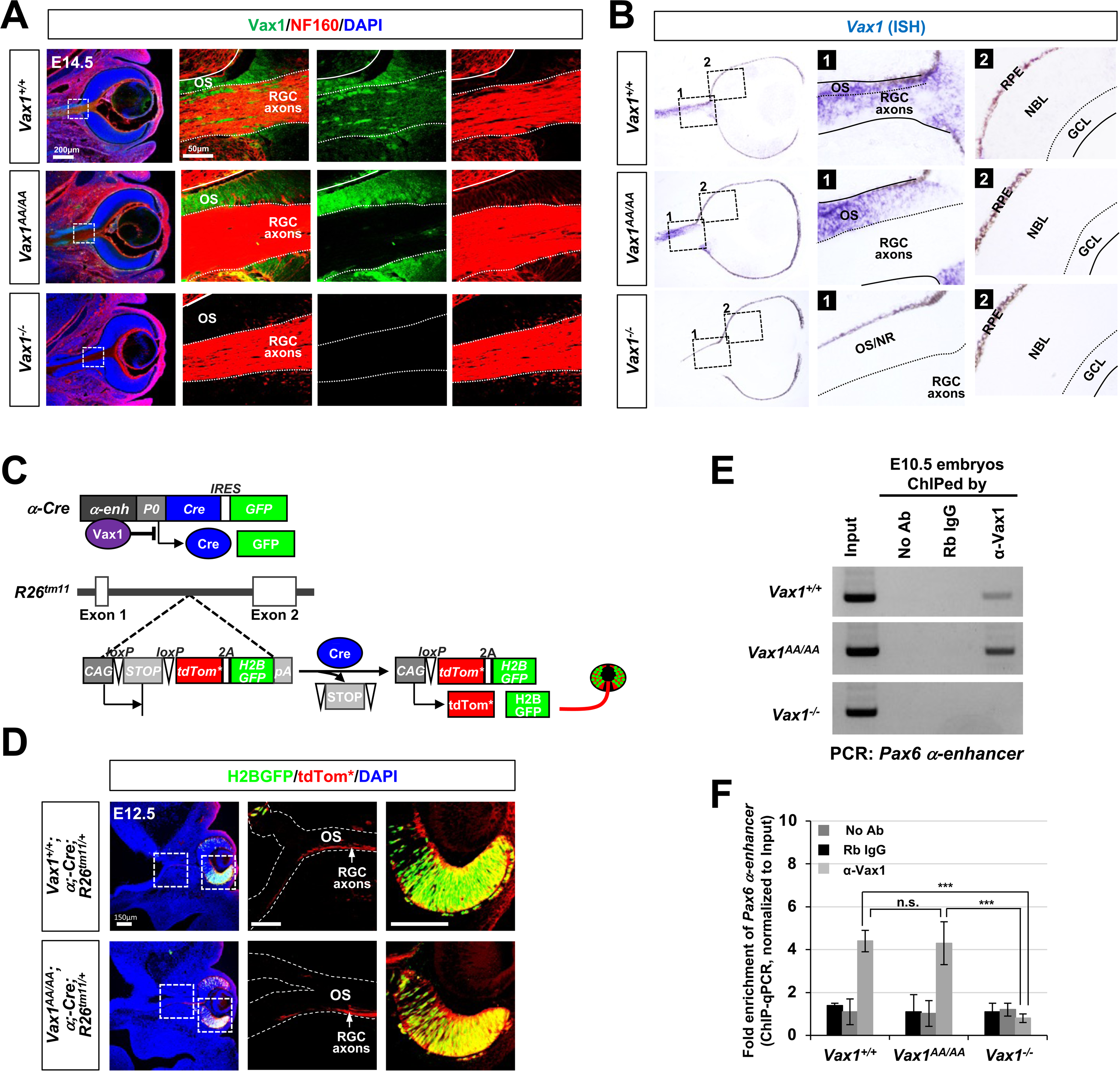
Vax1^AA^ exhibits defective intercellular transfer but intact transcription factor activity *in vivo*. **(A)** Intercellular transfer of Vax1 from E14.5 *Vax1^+/+^* and *Vax1^AA/AA^* littermate mouse OS cells to RGC axons was determined by immunostaining of the embryonic sections with α-Vax1 (green) and α-NF160 (red). **(B)** Expression of *Vax1* and *Vax1^AA^* mRNA in E14.5 *Vax1^+/+^* and *Vax1^AA/AA^* littermate mouse eyes and OS was examined by ISH. Boxed areas in the leftmost column are magnified in two right columns. The specificities of the anti-Vax1 antibody (A) and *Vax1* ISH probe (B) were determined by the absence of signals in E14.5 *Vax1^-/-^* mice. The *ISH* signals of the sense probes for *Vax1* did not exhibit specific signals (data not shown). The solid lines in the images indicate the boundary of OS, and the dotted-lines mark the border between the OS and RGC axon bundles. NR, neural retina; RPE, retinal pigment epithelium; NBL, neuroblast layer; GCL, ganglion cell layer. **(C)** The activities of a Vax1 target *Pax6 α*-enhancer in mouse embryos were measured by detecting H2B-EGFP, of which expression is induced together with membrane-bound tdTomato (tdTom*) at *R26^tm11^* gene locus after α-Cre- dependent excision of *loxP-STOP-loxP* cassette. Arrowheads point to the frontlines of the RGC axons. **(D)** Binding of Vax1 and Vax1^AA^ proteins on the *Pax6 α*-enhancer was determined by PCR detection of *Pax6 α*-enhancer sequences in DNA fragments isolated from E10.5 mouse embryonic cells by ChIP with indicated antibodies (see details in Methods). Input, mouse chromosomal DNA; No Ab, no antibody; Rb IgG, preimmune rabbit IgG; α-Vax1, rabbit anti-Vax1 polyclonal antibody. **(E)** Relative levels of *Pax6 α*- enhancer sequences in the ChIPed samples were compared by qPCR. The numbers in y- axis are average 2^-ΔCt^ values of the samples against critical threshold (Ct) values of Input. Error bars denote SD (n=5).

### Intact transcription factor activity of Vax1^AA/AA^ in vivo

We, next, examined whether the KA-to-AA mutation does not alter Vax1 transcription factor activity *in vivo,* as it did in cultured cells (Figure 1B), by monitored the expression of a reporter driven by *Pax6* α-enhancer, where Vax1 was identified to bind and suppress the enhancer activity (Mui et al., 2005). In practical, we used *α-Cre;R26^tm11^* mice (Marquardt et al., 2001; Wang et al., 2019), which express histone H2B-GFP (H2B-GFP) and membrane-bound tdTomato (tdTom*) reporters at *ROSA26* gene locus upon the excision of *loxP-STOP-loxP* (*LSL*) cassette by Cre recombinase expressed at downstream of *Pax6* α-enhancer (Figure 2C). The H2B-GFP signals were observed only in the retinal cells, but not in the OS cells, of E12.5 *Vax1^+/+^* mice, whereas the tdTom* signals were detectable in RGC axons in the OS as well as the retinal cells (Figure 2D, top row). The pattern was not changed in their *Vax1^AA/AA^* littermates (Figure 2D, bottom row). The results therefore suggest that *Pax6* α-enhancer activity was suppressed properly in the OS cells of *Vax1^+/+^* and *Vax1^AA/AA^* mice.

We also compared the abilities of Vax1 and Vax1^AA^ to bind *Pax6* α-enhancer sequences *in vivo*. Vax1-bound DNA fragments were isolated from E10.5 mouse heads by chromosome immunoprecipitation (ChIP) using an anti-Vax1 antibody and used for polymerase chain reaction (PCR) and quantitative PCR (qPCR) detection of *Pax6* α- enhancer sequences in the ChIPed DNA. The results revealed no significant difference in the binding abilities of Vax1 and Vax1^AA^ on the target sequences (Figure 2E and 2F). Together, these results suggest that the changes in *Vax1^AA/AA^* mice in comparing with *Vax1^+/+^* mice are unlikely resulted from the alteration of Vax1 target gene expression in the cells having active *Vax1* gene expression.

### Developmental delay of the OS in Vax1^AA/AA^ mice

Coloboma, the fissures in the ventral eyecup and OS, is observed in the eyes of humans and mice harboring *VAX1* mutations (Bertuzzi et al., 1999; Hallonet et al., 1999; Slavotinek et al., 2012). A coloboma was not observed in neonatal (i.e., post-natal day 0, P0) *Vax1^AA/AA^* mouse eyes unlike *Vax1^-/-^* mouse eyes (Figure 3A [middle row] and 3B; fissures in the colobomatous eyes are pointed by arrowheads). However, the optic fissures remained unclosed in *Vax1^AA/AA^* mice by E14.5, similar to *Vax1^-/-^* mouse eyes (Figure 3A [top row] and 3B). The unclosed OS in E14.5 *Vax1^AA/AA^* mice expressed Pax2 (paired homeobox 2), an OS marker, but not Pax6, a retinal marker, similar to the distribution observed in *Vax1^+/+^* littermates (Figure 3C [coronal], left and center columns). It contrasts with E14.5 *Vax1^-/-^* mice, which co-expressed Pax2 and Pax6 in the OS (Figure 3C [coronal], right column). Therefore, these results suggest that OS fate is specified properly in *Vax1^AA/AA^* mice, but not in *Vax1^-/-^* mice.

**Figure 3.**
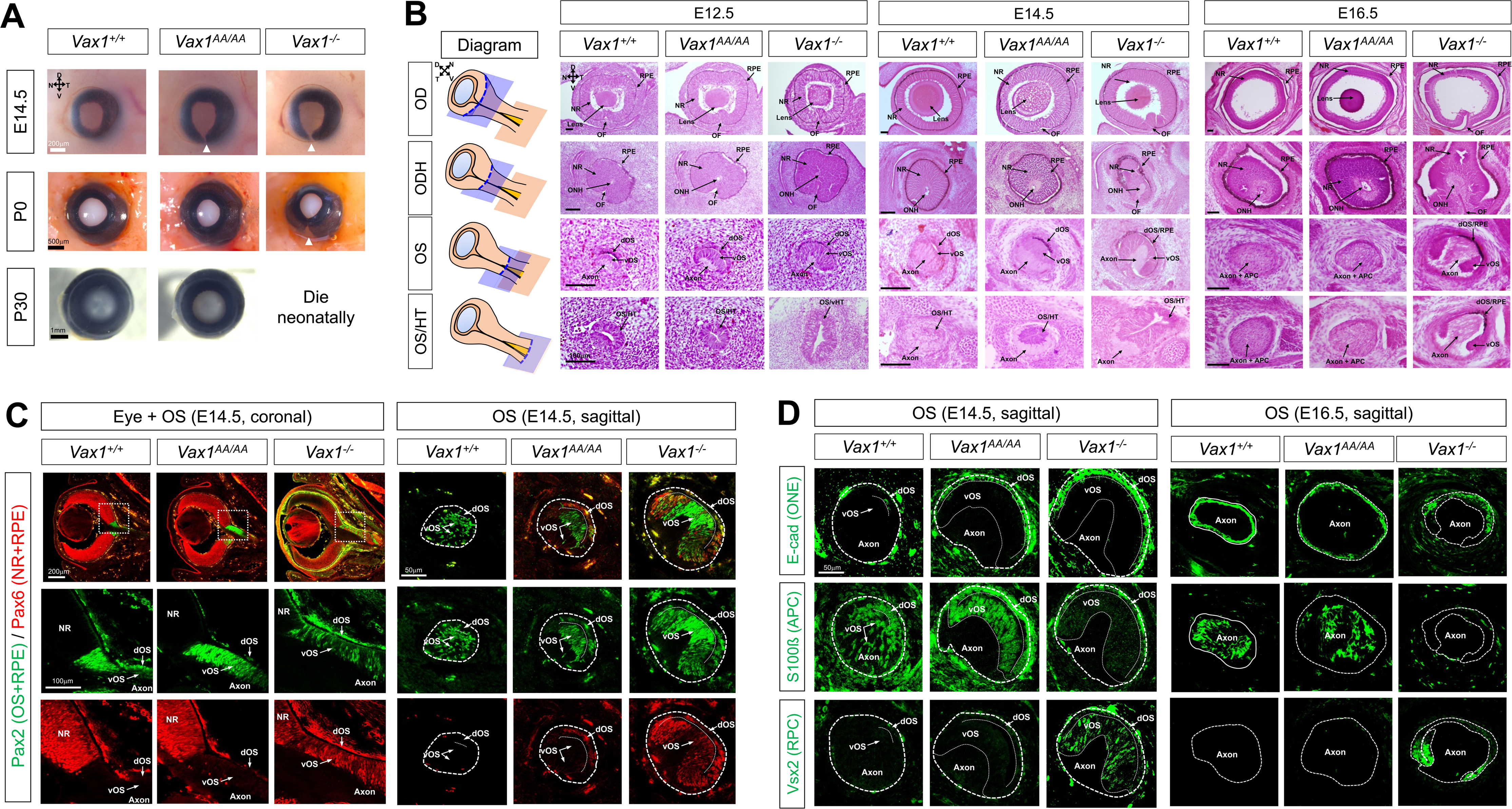
Developmental delay of OS differentiation in *Vax1^AA/AA^* mice. **(A)** Pictures of mouse eyes (frontal view) with the indicated genotypes are taken at various developmental stages. Arrowheads point to the optic fissures. N, nasal; T, temporal; D, dorsal; V, ventral. **(B)** Sagittal sections of mouse embryos were stained with hematoxylin & eosin (H&E). Positions of the sections are indicated by blue plates in the diagram (leftmost column). APC, astrocyte precursor cell; ODH, optic disc head; OF, optic fissure; dOS, dorsal optic stalk; vOS, NR, neural retina; RPE, retinal pigment epithelium; ventral optic stalk; vHT, ventral hypothalamus. Scale bars in the pictures are 100 μm. **(C)** Distributions of Pax6 and Pax2 in E14.5 mouse embryonic sections were examined by immunostaining. Boxed areas in top row images of coronal sections are magnified in two bottom rows. **(D)** Sagittal sections including medial OS of E14.5 and E16.5 mouse embryos were stained with the antibodies that recognize the corresponding markers. APC, astrocyte precursor cell; ONE, optic neuroepithelium; RPC, retinal progenitor cell.

The Pax2-positive OS cells were, however, clustered separately from RGC axons in E14.5 *Vax1^AA/AA^* mice, whereas they were spread among RGC axons in E14.5 *Vax1^+/+^* mice (Figure 3C, sagittal). The OS cells in E14.5 *Vax1^+/+^* mice expressed an astrocyte precursor cell (APC) marker, S100 calcium-binding protein ß (S100ß), but had lost a neuroepithelium marker, E-cadherin (E-cad) (Figure 3D, left column of E14.5 panel). The cells in the ventral OS (vOS) of E14.5 *Vax1^AA/AA^* mice also expressed S100 without E-cad, while those in the dOS co-expressed S100ß and E-cad (Figure 3D, center column of E14.5 panel). The vOS cells in E14.5 *Vax1^-/-^* mice, however, did not express S100ß, instead they expressed the retinal marker Vsx2 (Figure 3D, right column of E14.5 panel). In E16.5 *Vax1^AA/AA^* mice, similar to the distribution observed in *Vax1^+/+^* littermates, S100ß- positive OS cells had entirely lost E-cad expression and started to disperse between RGC axons, whereas S100ß-negative OS cells still formed Ecad-positive neuroepithelial clusters in E16.5 *Vax1^-/-^* mice (Figure 3D, E16.5 panel). These results suggest that differentiation of S100ß-positive APC from the OS neuroepithelium is impaired in *Vax1^-/-^* mice but merely delayed in *Vax1^AA/AA^* mice.

### Growth retardation of RGC axons in the OS of Vax1^AA/AA^ mice

We next examined whether RGC axons could grow in *Vax1^AA/AA^* mouse OS as fast as those in *Vax1^+/+^* mouse OS. To visualize RGC axons in 3-dimension (3D), we reconstituted tdTom* fluorescence reporter signals of tissue-cleared *Vax1^+/+^;α-Cre;R26^tm11^* and *Vax1^AA/AA^;α-Cre;R26^tm11^* mouse embryos after the lightsheet microscopic imaging (Figure 4A; see details in Methods). In E12.5, the RGC axons are observed commonly in the OS of *Vax1^+/+^* and *Vax1^AA/AA^* littermate mice (Figure 4B [left panel]; Video 1 and 2), implicating that RGC axons exit from the eyes properly in *Vax1^AA/AA^* mice. However, the lengths of RGC axons in E12.5 *Vax1^AA/AA^* mice are shorter than that of *Vax1^+/+^* littermates (Figure 4C). The RGC axon terminals were still observed in the OS of E13.5 *Vax1^AA/AA^* mice, while those had passed the OS/HT borders and reached the vHT midline in *Vax1^+/+^* littermates (Figure 4B [center panel] and 4C; Video 3 and 4). The length of RGC axons in E14.5 *Vax1^AA/AA^* mice remained shorter than that of *Vax1^+/+^* littermate mice (Figure 4B [right panel]; Video 5 and 6). Moreover, RGC axons were absent in the vHT midline but found in the lateral HT wall in E14.5 *Vax1^AA/AA^* mice, whereas the axons extended underneath the vHT to form the OC in E14.5 *Vax1^+/+^* littermate. The numbers of Brn3b- positive RGCs in E14.5 *Vax1^+/+^*and *Vax1^AA/AA^* mouse retinas were not different significantly each other (Figure 4 – figure supplement 1), suggesting that the weak axon signals in *Vax1^AA/AA^* mouse OS were not resulted from the decrease of RGC numbers. Together, these results suggest that RGC axons grow slowly in the OS to arrive at the HT late and do not access vHT midline but extend ipsilaterally in *Vax1^AA/AA^* mice.

**Figure 4.**
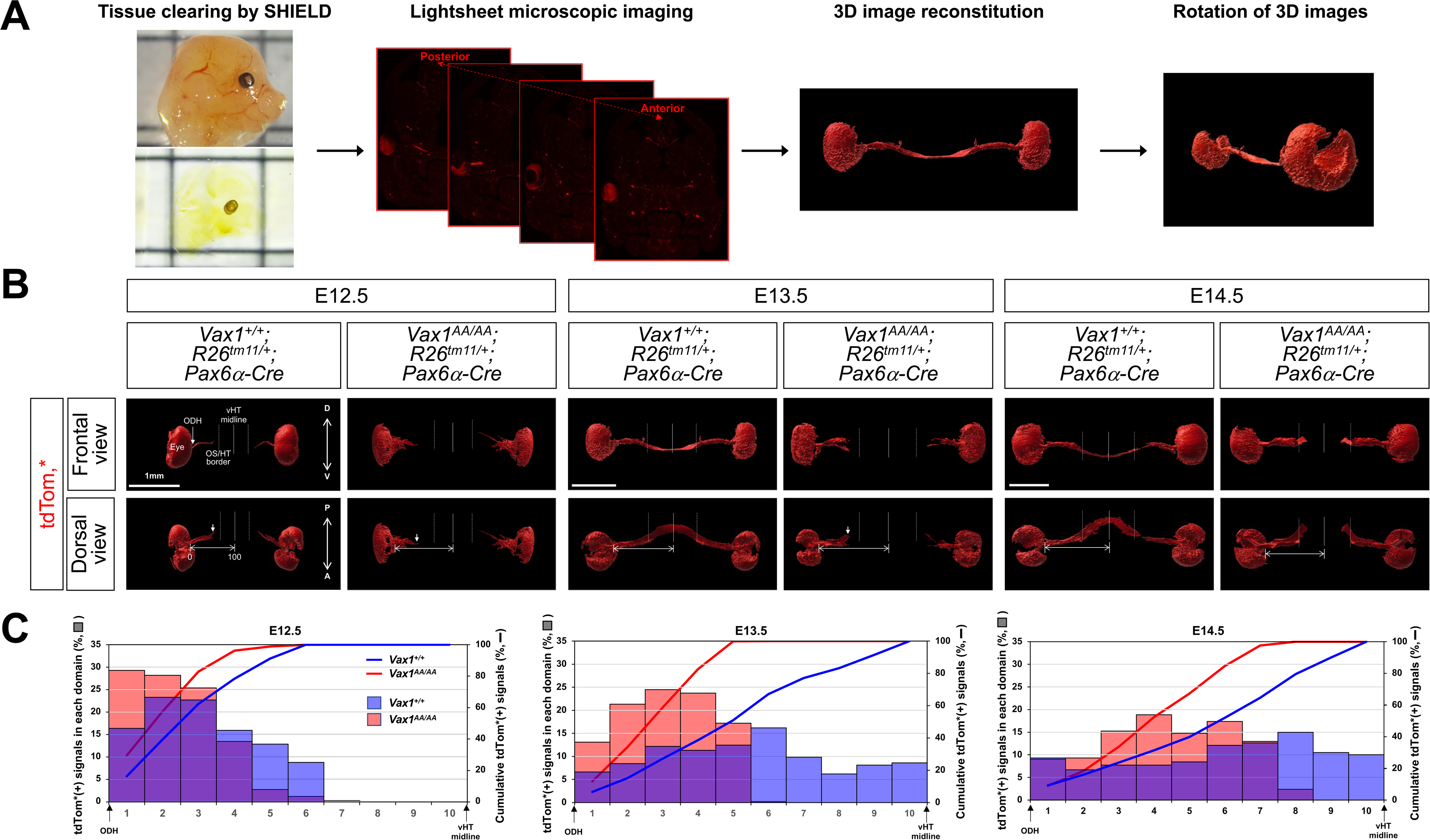
Retarded RGC axon growth in *Vax1^AA/AA^* mice. **(A)** Schematic diagram depicts 3D reconstitution of tdTom* fluorescent signals in RGC axons. Retinal cells, including RGCs, in *Vax1^+/+^* and *Vax1^AA/AA^* littermate mouse embryos, which were cleared by the SHIELD method, were visualized by tdTom* reporter expressed in *R26^tm11^* gene locus in Pax6 α-Cre-dependent manner. The tdTom* signals of retinal cells and RGC axons, which grow in the OS to form the OC prior to extending through the OT, were then reconstituted in 3D (see details in Methods). **(B)** The 3D reconstituted images of E12.5, E13.5, and E14.5 mouse embryos are shown. **(C)** Relative tdTom* intensities in each domain of RGC axon track, which is divided into 10 segments from optic nerve head (OHD) to vHT midline, are shown in the graphs. Cumulative tdTom* intensities, which represent the positions of RGC axon terminals, are also shown in the graphs. The values are averages (n=3; 3 independent litters).

Despite the growth retardation of RGC axons, *Netrin-1*, a RGC axon growth factor (Deiner and Sretavan, 1999), was found in the OS of *Vax1^AA/AA^* mice, whereas it was disappeared in *Vax1^-/-^* mouse OS (Figure 2 - figure supplement 2A and 2B). The expression of Semaphorin-5A (Sema5A), which is a repulsive RGC guidance cue expressed in the OS cells (Oster et al., 2003), was also expressed properly in the OS of *Vax1^+/+^* and *Vax1^AA/AA^* mice, but not in *Vax1^-/-^* mouse OS (Figure 4 - figure supplement 2C and 2D). The results suggest that retarded RGC axon growth in *Vax1^AA/AA^* mouse OS was not resulted from the altered expression of these RGC axon guidance cues.

### Structural alteration of the vHT in achiasmatic Vax1^AA/AA^ mouse embryos

We found the HT in *Vax1^AA/AA^* and *Vax1^-/-^* mouse embryos protruded ventrally without underlying RGC axons bundles, while the vHT of E14.5 *Vax1^+/+^* mice was flattened over RGC axons forming the OC (Figure 5A and 5B). However, *Shh*, which is critical for specification of the HT and OS (Chiang et al., 1996; Kim and Lemke, 2006), showed a similar expression pattern in E14.5 *Vax1^+/+^*, *Vax1^AA/AA^* and *Vax1^-/-^* mice (Figure 5C, top row). This suggests that fate specification of the vHT is likely unaffected in *Vax1^AA/AA^* and *Vax1^-/-^* mice, as it had been shown in previous reports (Bertuzzi et al., 1999; Hallonet et al., 1999). The Glast-positive radial glia, which express various RGC axon guidance cues (Herrera et al., 2019; Petros et al., 2008), were also observed in the vHT of *Vax1^AA/AA^* and *Vax1^-/-^* mice, indicating normal development of vHT radial glia from Nestin-positive neuroepithelium in the mice (Figure 5C, second and third rows).

**Figure 5.**
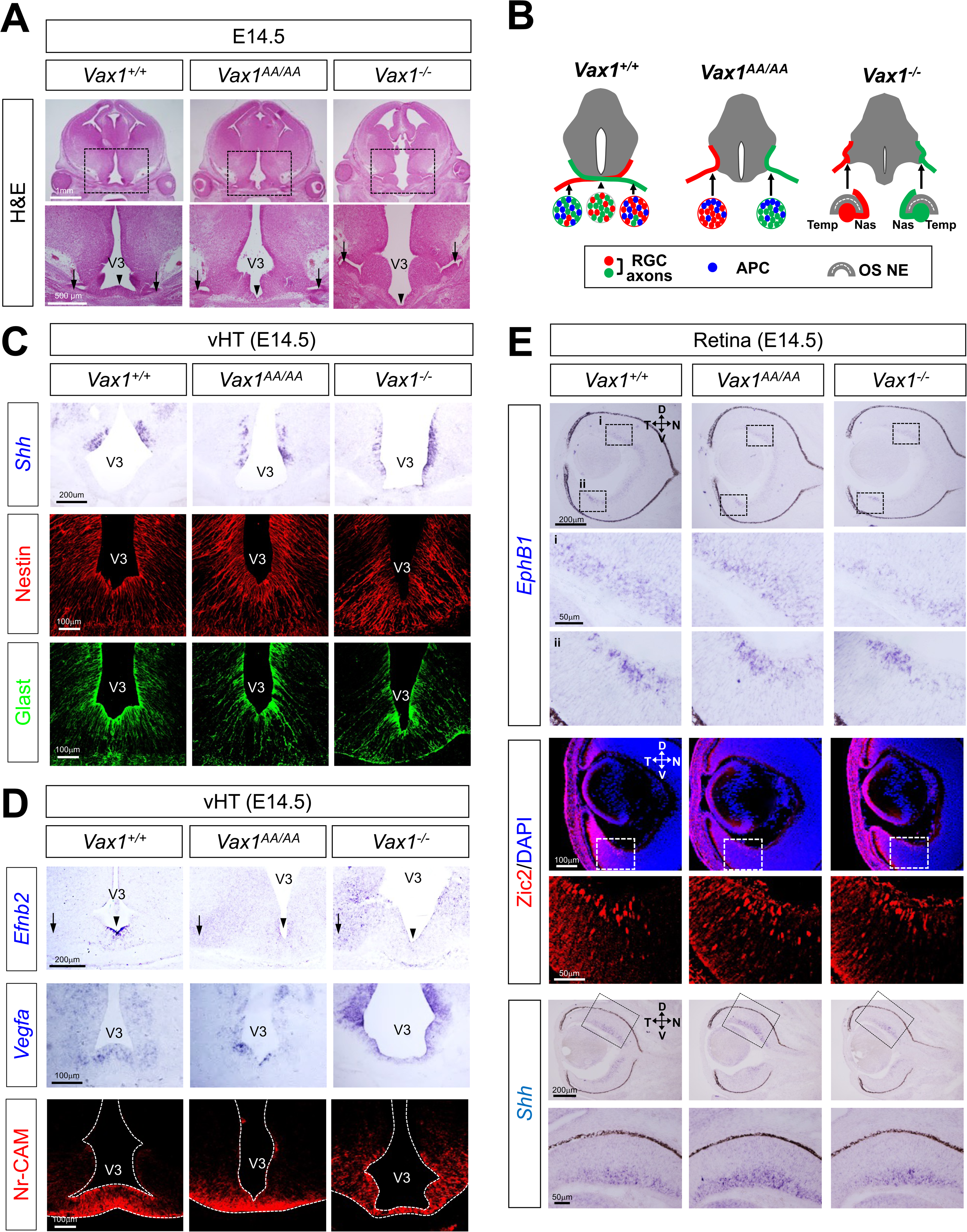
Expression of RGC axon guidance cues in *Vax1^AA/AA^* mice. (**A**) Coronal sections of E14.5 mouse embryos were stained with H&E. Arrows indicate OS/HT junction and arrowheads point vHT midline. V3, third ventricle. **(B)** Schematic diagrams that show the structures of OS/HT junctions and RGC axon pathways in *Vax1^+/+^*, *Vax1^AA/AA^*, and *Vax1^-/-^* mice. Nas, nasal; Temp, temporal. OS NE, OS neuroepithelium. **(C)** Distributions of *Shh* mRNA in the embryonic sections were detected by *in situ* hybridization (ISH). Development of hypothalamic radial glia (RG) from NPC was determined by immunostaining for an RG marker Glast (glutamate aspartate transporter 1) and an NPC marker Nestin, respectively. **(D)** Expression of a repulsive guidance cue, ephrinB2, and the attractive guidance cues, Vegfa and Nr-CAM, for RGC axons in E14.5 mouse vHT was examined by *ISH* (*Efnb2* and *Vegfa*) and IHC (Nr-CAM). Arrows indicate OS/HT junction and arrowheads point vHT midline. **(E)** Expressions of retinal genes that induce ipsilateral RGC axon projection were examined by ISH (*EphB1* and *Shh*) and IHC (Zic2). Boxed areas in top row images are magnified in bottom rows. The *ISH* signals of the sense probes for *Efnb2*, *Vegfa*, *Ephb1*, and *Shh* did not exhibit specific signals (data not shown).

We, thus, examined whether the altered expression of RGC axon guidance cues in the vHT is related with the failure of RGC axon growth toward the vHT midline in *Vax1^AA/AA^* and *Vax1^-/-^* mice. However, attractive guidance cues for RGC axons, including Vegfa and Nr-Cam (Erskine et al., 2011; Williams et al., 2006), were expressed properly in the vHT of E14.5 *Vax1^AA/AA^* and *Vax1^-/-^* mice (Figure 5D, second and third rows). Despite the failure of RGC axonal projection to the midline (Figure 4B), the expression of *Ephrin-B2* (*Efnb2*), which triggers the repulsion of EphB1-expressing RGC axons from the VT mouse retina (Williams et al., 2003), was rather decreased in the vHT midline cells of E14.5 *Vax1^AA/AA^* and *Vax1^-/-^* mice in comparison to that in E14.5 *Vax1^+/+^* littermates’ vHT midline glia (Figure 5D, arrows in top row).

Instead, *Efnb2* mRNA was detectable in the lateral HT area in E14.5 *Vax1^AA/AA^* and *Vax1^-/-^* mice at a relatively lower level than that in the midline glia of *Vax1^+/+^* mice (Figure 5D, arrowheads in top row). Given no repulsion of the majority RGC axons, except for those from the VT retina, by high level Ephrin-B2 in the vHT midline glia in *Vax1^+/+^* mice (Williams et al., 2003), the low level Ephrin-B2 in the lateral HT of *Vax1^AA/AA^* mice should be insufficient to trigger repulsive responses in their RGC axons. The expressions of *EphB1* and its upstream regulator Zic2 were enriched in the VT RGCs of E14.5 *Vax1^AA/AA^* and *Vax1^-/-^* mouse retinas, similar to the expression patterns in E14.5 *Vax1^+/+^* mice (Figure 5E) (Herrera et al., 2003). These results implicate that there might be no ectopic activation of EphB1 in *Vax1^AA/AA^* and *Vax1^-/-^* mouse RGC axons. Shh, which was identified as a transacting repulsive cue in mouse RGC axons (Peng et al., 2018), was also expressed properly in RGCs of those mice (Figure 5E). Together, our data suggest that *Vax1^AA/AA^* mouse RGC axons might be repelled from the lateral HT by other signals than Ephrin-B2 and Shh. It is also possible that RGC axons in *Vax1^AA/AA^* mice could not access to the vHT midline by losing axon growth signals other than Vegfa and Nr-Cam.

### Ipsilaterally biased retina-brain connections in the Vax1^AA^ mice

Using adult *Vax1^AA/AA^* mice, we were able to analyze the structures and functions of the mouse visual system, which are difficult to study in *Vax1^-/-^* mice because they die perinatally (Bertuzzi et al., 1999; Hallonet et al., 1999). We found no significant difference in brain size and shape between P30 *Vax1^+/+^* and *Vax1^AA/AA^* littermate mice (Figure 6A and 6C [left column]). The olfactory bulb (OB) of *Vax1^AA/AA^* mice also appeared normal (Figure 6C, the image in left bottom corner), although a previous report noted hypoplastic OBs in the few surviving *Vax1^-/-^* mice (Soria et al., 2004). Furthermore, *Vax1^AA/AA^* mice have only one pituitary gland (data not shown), whereas *Vax1^-/-^* mice have an extra pituitary gland by virtue of failure to suppress ectopic pituitary fate specification in the ventral anterior forebrain (Bharti et al., 2011). These results also suggest that Vax1-dependent specification of ventral forebrain structures is not significantly altered in *Vax1^AA/AA^* mice.

**Figure 6.**
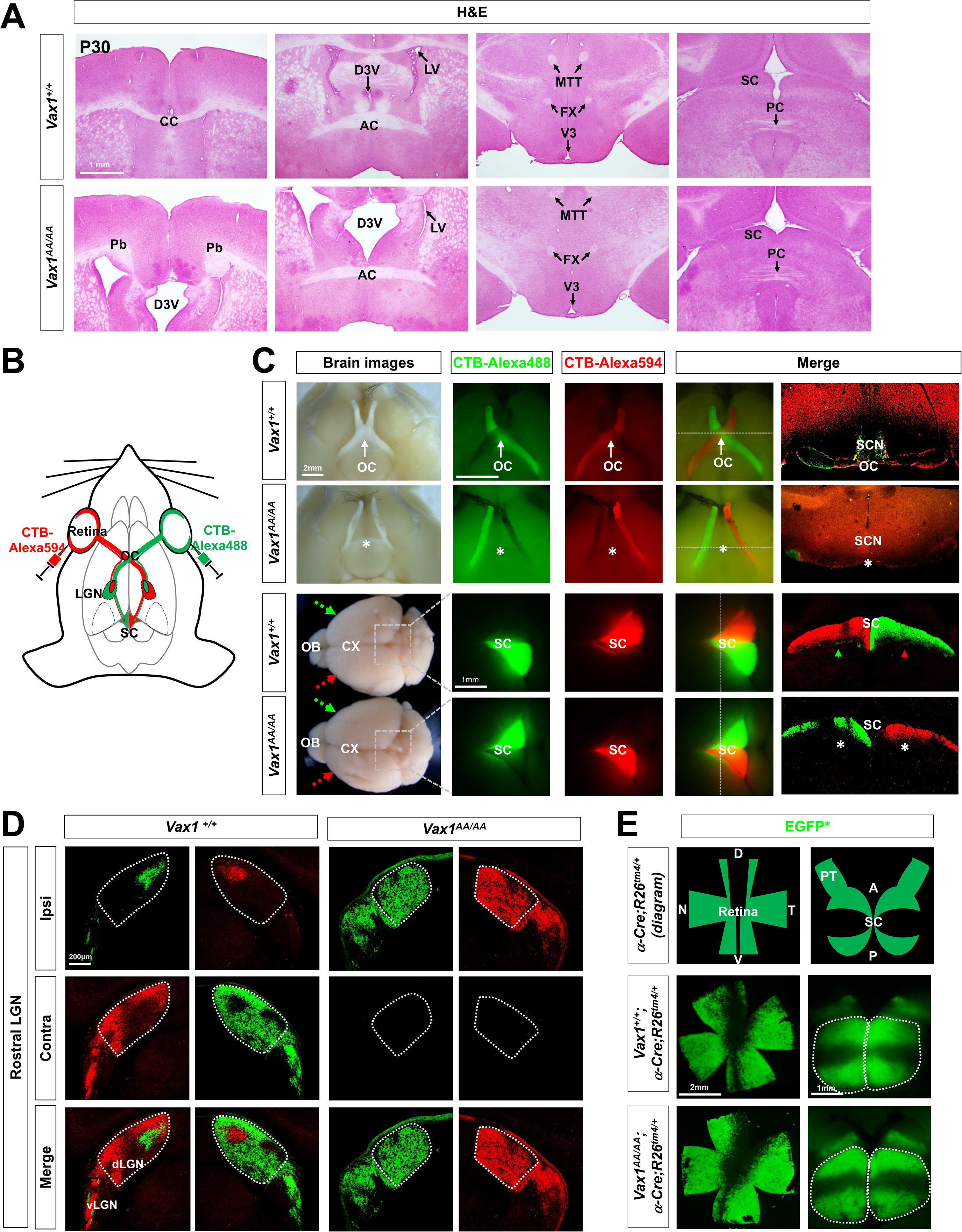
Ipsilaterally-biased RGC axon projection in *Vax1^AA/AA^* mice. **(A)** Coronal sections of P30 mouse brain were stained with H&E. The sections containing the indicated commissural structures were identified and shown. AC, anterior commissure; CC, corpus callosum; D3V, dorsal third ventricle; FX, fornix Pb, probus bundle; PC, posterior commissure; SC, superior colliculus; MTT, mammillo-thalamic tract. **(B)** Diagram depicts fluorescence labeling of mouse RGCs. **(C)** Alexa488- and Alexa594-labeld CTB proteins were injected into right and left eyes of the mice at P28, respectively. Brains of the CTB- injected mice were isolated at P30 and fluorescent signals emitted from the brains were visualized by Axio Zoom stereoscope (Zeiss; three center columns), and then the fluorescent signals in coronal sections of the brains were detected by FV1000 confocal microscope (Olympus; rightmost column). Bright-field images of the brains were also taken to show the structure of the brains and optic nerves (leftmost column). CX, cerebral cortex; OB, olfactory bulb; *, ventral midline of HT equivalent to the position of OC in *Vax1^+/+^* littermates. **(D)** Coronal sections of the CTB-injected mouse brains were collected and fluorescent signals in the LGN area were detected. The areas surrounded by dotted lines are dLGN. Contra, contralateral LGN section; Ipsi, ipsilateral LGN section. **(E)** The retinas and brains of P30 *Vax1^+/+^* and *Vax1^AA/AA^* littermate mice carrying α-*Cre*;*R26^tm4/+^* transgenes, which express membrane-targeted EGFP (EGFP*) at *R26^tm4^* gene locus upon Cre-dependent excision of *loxP-tdTomato*-loxP* gene cassette, were isolated and visualized. The areas surrounded by dotted lines are SC. PT, pretectum.

Among the commissures that are reported to be absent in *Vax1^-/-^* mice (Bertuzzi et al., 1999), the CC and OC are missing in *Vax1^AA/AA^* mice, whereas the AC and fornix are present (Figure 6A and 6C [left column]). The posterior commissure (PC) was also present and appropriately positioned beneath the SC in *Vax1^AA/AA^* mice (Figure 6A, rightmost column). Further, we visualized the RGC axons of mice using fluorescent dye-conjugated cholera toxin B (CTB) protein (Luppi et al., 1990). The Alexa Fluor 488-conjugated CTB and Alexa Fluor 594-conjugated CTB were injected into the right and left eyes, respectively, of *Vax1^+/+^* and *Vax1^AA/AA^* littermate mice at postnatal day 28 (P28; Figure 6B). Fluorescence signals emitted by CTB-labeled RGC axons in the mice were then detected at P30 (Figure 6C). Green and red fluorescence signals were predominantly detected in RGC axons that projected across the midline in *Vax1^+/+^* mice (Figure 6C, top row). Consequently, the majority of CTB fluorescence signals were observed in the contralateral SCs of *Vax1^+/+^* mice (Figure 6C, third row). In contrast, fluorescence signals of CTB were detected exclusively in the ipsilateral SCs of *Vax1^AA/AA^* mice, in which it was not possible to detect the OC where differentially labeled RGC axons met (Figure 6C, second and bottom rows).

We also examined whether ipsilaterally projecting *Vax1^AA/AA^* mouse RGC axons were properly connected to the dLGN, a thalamic nucleus that relays visual information from the retina to the visual cortex (Seabrook et al., 2017). Axons from RGCs in the ventral-temporal mouse retina project to the ipsilateral dLGN, where the majority of RGC axons from the contralateral retina are connected (Herrera et al., 2019; Petros et al., 2008). The minor ipsilateral axon terminals are then segregated from the major contralateral axon terminals by a retinal activity-dependent refinement process during postnatal days (Guido, 2018; Huberman et al., 2008). Segregation of binocular RGC axons was clearly seen in the dLGN of P30 *Vax1^+/+^* mice (Figure 6D, left two columns). However, RGC axons in the dLGN of P30 *Vax1^AA/AA^* mice originated only from the ipsilateral retina, and no contralateral RGC axons were observed in the dLGN (Figure 6D, right two columns).

We further examined whether retinocollicular topographic connectivity was established properly in the *Vax1^AA/AA^* mouse SC, which lacks contralateral RGC axon terminals (Figure 6C, bottom row). Axons from RGCs in the temporal retina are known to connect to the anterior SC, whereas those from the nasal retina link to the posterior SC (Lemke and Reber, 2005). In the perpendicular axis, RGC axons from the dorsal retina arrive at the lateral SC, and those from the ventral retina are wired to the medial SC. Thus, axons of Pax6 α-Cre-affected RGCs in the ventral and peripheral retina, which express membrane-localized EGFP (EGFP*) in *ROSA26* gene locus of *R26^tm4^* Cre reporter mice (Marquardt et al., 2001; Muzumdar et al., 2007), map to medial and peripheral SC areas in P30 *Vax1^+/+^* mice (Figure 6E, middle row). This pattern was also observed in the SCs of *Vax1^AA/AA^* littermate mice (Figure 6E, bottom row), implying that the retinocollicular topography is established properly in *Vax1^AA/AA^* mice.

### Light-stimulated retinal and cortical responses in Vax1^AA/AA^ mice

Next, we examined whether visual information can be processed in the retina and delivered to the brain in *Vax1^AA/AA^*mice. First, we tested the activities of mouse retinas using electroretinography (ERG) recordings. The shapes of ERG a-waves of dark-adapted P45 mouse retinas, which reflect the function of rod photoreceptors, appeared normal in *Vax1^AA/AA^* mice, although the amplitudes of ERG b-waves, which are generated by bipolar cells and Müller glia in the inner retina downstream of photoreceptors (Miura et al., 2009), were decreased slightly (Figure 7A [right column] and 7B). On the contrary, the amplitudes of ERG a-waves of light-adapted mouse retinas, which reflect the activity of cone photoreceptors, were decreased significantly in *Vax1^AA/AA^* mice compared with *Vax1^+/+^* littermate mice (Figure 7A [left column] and 7C [graph in left]), leading to a consequent decrease in amplitudes of ERG b-waves (Figure 7A [left column] and 7C [graph in right]). The reduced photopic ERG responses in *Vax1^AA/AA^* mice were likely related to the decrease in S-opsin–positive cone photoreceptors in the ventral retina of *Vax1^AA/AA^* mice, which did not show no significant differences in other cell types in the retina and optic nerves (Figure 7 - figure supplement 1).

**Figure 7.**
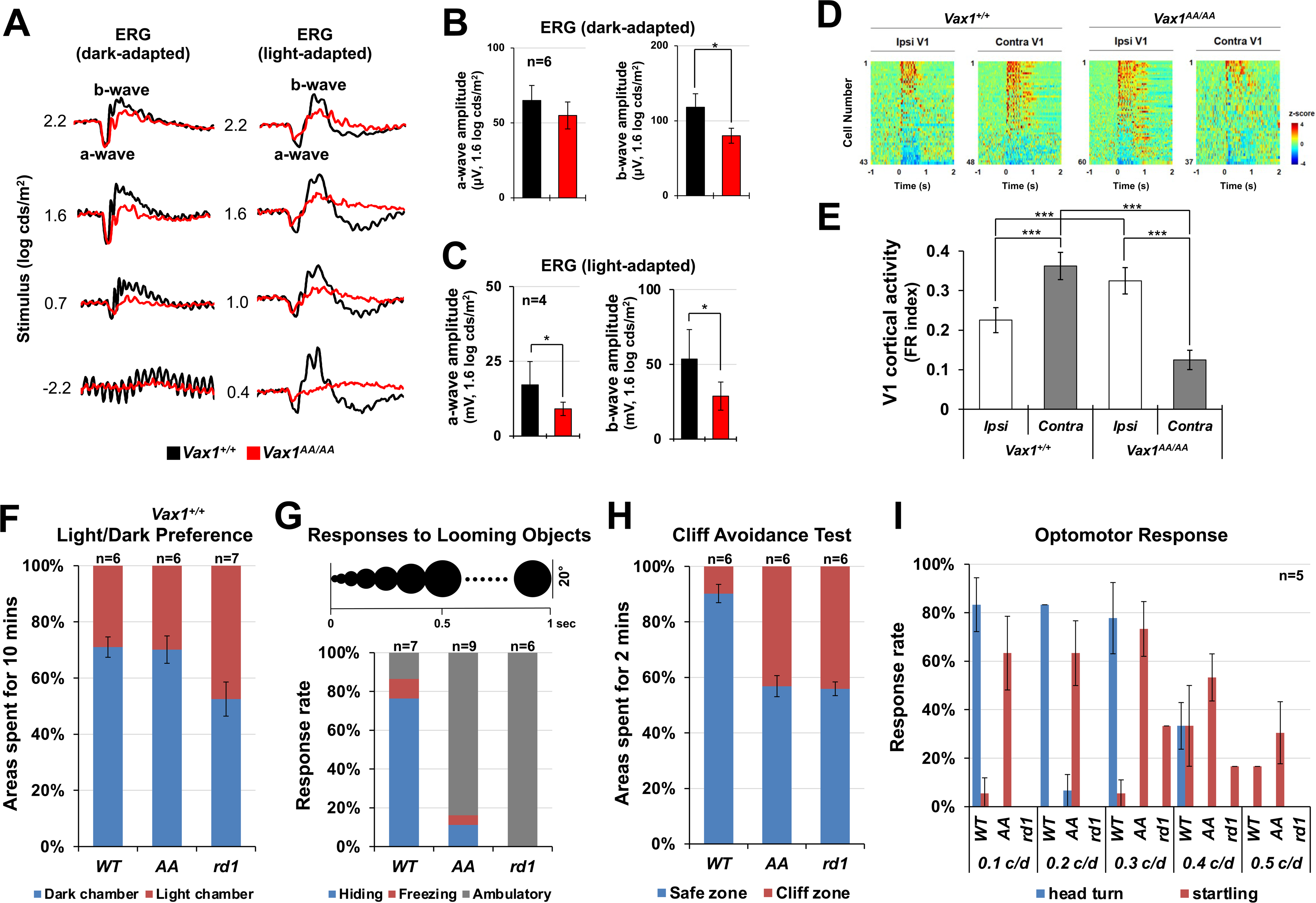
Reduced visual acuity of *Vax1^AA/AA^* mice. **(A)** Electrophysiological activities of P45 *Vax1^+/+^* and *Vax1^AA/AA^* mouse retinas were examined by ERG (see details in Methods). The amplitudes of scotopic **(B)** and photopic **(C)** ERG a- and b-waves at 1.6 log cds/m^2^ condition are presented. Numbers of mice tested are given in the graphs (4 independent litters). **(D)** Light-evoked excitation of cortical neurons in P45 *Vax1^+/+^* and *Vax1^AA/AA^* mouse V1 were measured by silicon multielectrode probes after monocular illumination (see details in Methods). Responses of neurons in the monocular zones of ipsilateral (Ipsi) and contralateral (Contra) visual cortices were recorded. Color-coded heatmap represents average z-scores of spike firing rates. Red indicates increase and blue indicates decrease of firing rates from the baseline, respectively. **(E)** Mean firing-rate change index of ipsilateral and contralateral V1 neurons in *Vax1^+/+^* and *Vax1^AA/AA^* mice. y-axis values are mean ± SEM. **(F)** Relative occupancy of light and dark chambers by P45 *Vax1^+/+^* (*WT*), *Vax1^AA/AA^* (*AA*), and *Pde6b^rd1/rd1^* (*rd1*) mice for 10-min measurement period was determined and shown in a graph. **(G)** Escaping responses of the mice, which were given the expanding black circle on top screens to mimic a looming shadow of a predator, were determined and shown in a graph. **(H)** To determine stereoscopic vision of the mice, relative occupancy of safe and cliff zones by the mice for 2-min measurement period was determined and shown in a graph. **(I)** Accuracy of the mice to turn their heads to the directions where black and white vertical stripes rotate was measured and shown in a graph (head turn). Startling responses of the mice in response to the stimuli were also measured (startling). Numbers of the mice tested are given in the graphs in F – H.

We also examined whether visual information received by the retina is delivered to the visual cortex via the dLGN in *Vax1^AA/AA^* mice. To this end, we recorded electrical activities of neurons in the monocular zone of the primary visual cortices (V1) in both hemispheres of mouse brain while giving the visual stimuli only to the left eye. We found that it predominantly triggered responses of cortical neurons in the right (i.e., contralateral) V1 of *Vax1^+/+^* mice; conversely, it mainly activated neurons in the left (i.e., ipsilateral) V1 of *Vax1^AA/AA^*mice (Figure 7D and 7E). These results demonstrate that *Vax1^AA/AA^* mouse retina send the signals to the ipsilateral dLGN for subsequent delivery of the information to the visual cortex, whereas the retinal signals are sent mainly to the contralateral routes in *Vax1^+/+^* mice.

### Reduced visual acuity of Vax1^AA/AA^ mice

We next examined whether the ipsilaterally-biased retinogeniculate and retinocollicular pathways influenced the visual responses of *Vax1^AA/AA^* mice. First, we tested whether the mice could discriminate light and dark spaces. We found that P45 *Vax1^+/+^* and *Vax1^AA/AA^* littermate mice spent longer periods in the dark chamber than in the illuminated chamber (Figure 7F). On the contrary, P45 retinal dystrophy 1 (*rd1*) mutant *C3H/HeJ* blind mice, which have a homozygous mutation of *phosphodiesterase 6b* (*Pde6b; Pde6b^rd1/rd1^)* gene (Chang et al., 2002), spent equivalent periods in dark and light chambers (Figure 7F). These results suggest that *Vax1^AA/AA^* mice discriminate light and dark space as efficiently as *Vax1^+/+^* mice.

Second, we examined whether mice could recognize images that mimic a looming shadow of predator (Yilmaz and Meister, 2013). P45 mice were placed in a transparent box covered by a computer monitor displaying a black circle that expands by a 20° angle from the mouse head. *Vax1^+/+^* mice froze or hid under a shelter while the circle in the monitor expanded (Figure 7G; Video 7). *Vax1^AA/AA^* mice also exhibited hiding and/or freezing responses; however, their response frequency was significantly lower than that of *Vax1^+/+^* mice (Figure 7G; Video 8). *Pde6b^rd1/rd1^* mice showed no behavioral responses at all (Figure 3D; Video 9). These results suggest that *Vax1^AA/AA^* mice could recognize the shadow pattern, but did so less efficiently than *Vax1^+/+^* mice.

Third, we investigated whether mice could discriminate near and far objects by placing them on a transparent plate, half of which was printed with a flannel pattern image (i.e., safe zone) and the other unprinted half was extended over a floor printed with the same flannel pattern (i.e., cliff zone) (Sloane et al., 1978). P45 *Vax1^+/+^* mice stayed in the safe zone and rarely ventured into the cliff zone (Figure 7H; Video 10). However, *Vax1^AA/AA^* littermate mice moved freely between cliff and safe zones (Figure 7H; Video 11), as did P45 *Pde6b^rd1/rd1^* blind mice (Figure 7H; Video 12). These results suggest that *Vax1^AA/AA^* mice might have impaired depth perception.

Last, we also assessed the visual acuity of mice by measuring optomotor responses (OMRs) to horizontally moving black and white vertical strips using the OptoMotry (Prusky et al., 2004). P45 *Vax1^+/+^* mice turned their heads in the direction of vertical stripe movement (Figure 7I; Video 13); however, P45 *Vax1^AA/AA^* and *Pde6b^rd1/rd1^* mice failed to show a valid OMR (Figure 7I; Video 14 and 15). These results suggest that the visual acuity of *Vax1^AA/AA^* mice is significantly compromised compared with that of *Vax1^+/+^* mice.

### Seesaw nystagmus of Vax1^AA/AA^ mice

The OMR visual acuity test counts head-turn events to moving objects. Therefore, mice may not respond properly if their oculomotor system is abnormal. Interestingly, *Vax1^AA/AA^* mice startled in response to the stimuli, although they did not show corresponding head-turn behaviors (Figure 7I; Video 14). This contrasted with *Pde6b^rd1/rd1^* blind mice, which showed no stimulus-dependent responses (Figure 7I; Video 15). These results suggest that the reduced visual acuity of *Vax1^AA/AA^* mice might have resulted from an abnormal visuomotor system that cannot trigger proper head-turn responses.

Moreover, achiasmatic humans and dogs were also reported to have reduced visual acuity (Apkarian et al., 1995; Dell’Osso and Williams, 1995; Williams et al., 1994). Interestingly, they commonly exhibited seesaw nystagmus—an out of phase vertical oscillation in the two eyes in the absence of a visual stimulus. We found the eyes of P45 *Vax1^AA/AA^* mice oscillated spontaneously in an oval track in the absence of visual stimulus (Figure 8A and 8B; Video 17), whereas the eyes of *Vax1^+/+^* littermate mice gazed stably (Figure 8A, left column; Video 16). The oscillatory cycles of the two eyes of *Vax1^AA/AA^* mice were out of phase in the vertical axis but in phase in the horizontal axis (Figure 8A, right column; Figure 8 - figure supplement 1), phenocopying the seesaw nystagmus of achiasmatic humans and dogs.

**Figure 8.**
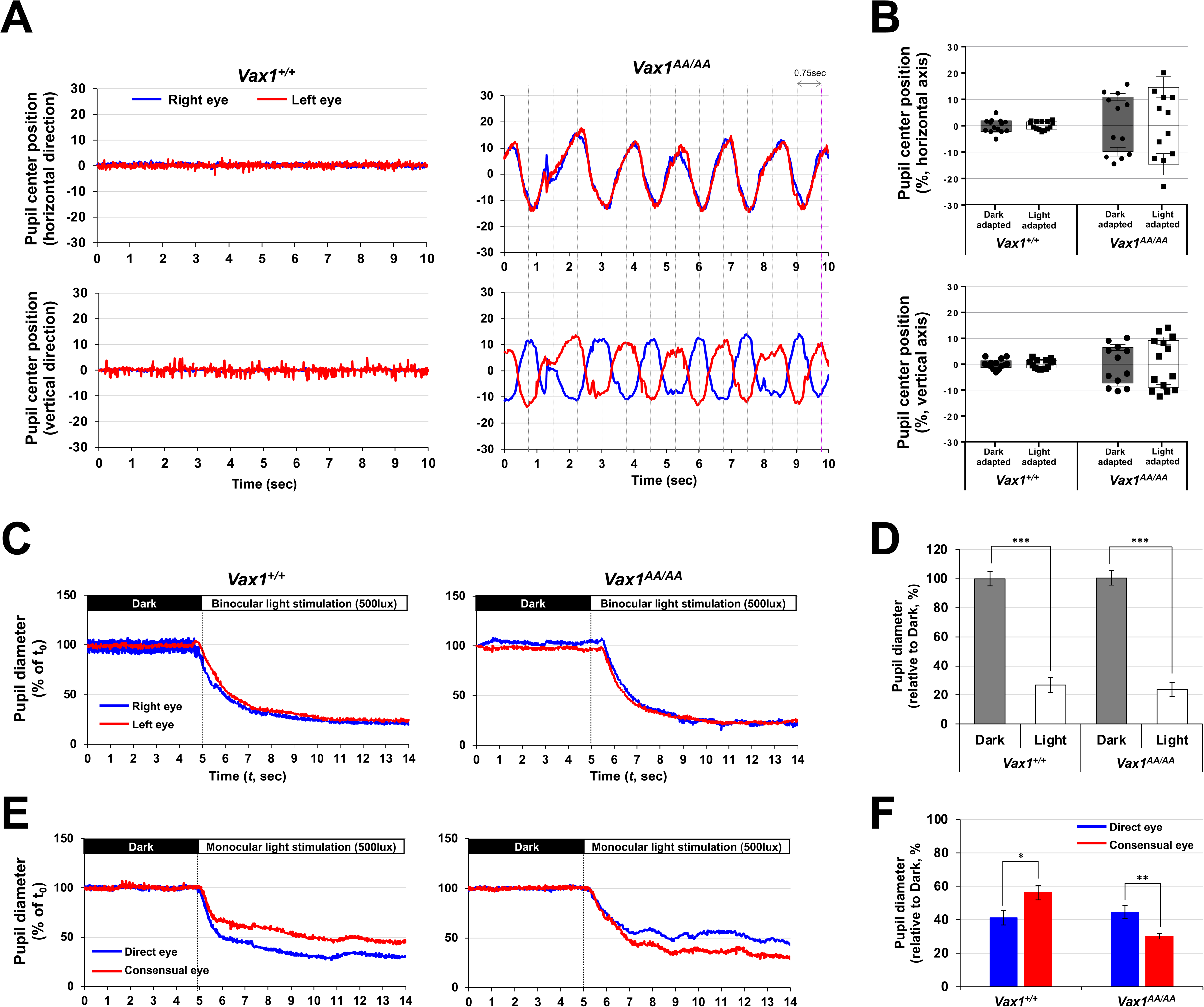
Seesaw nystagmus of *Vax1^AA/AA^* mice. **(A)** Positions of pupil centers in right and left eyes of head-fixed P45 *Vax1^+/+^* and *Vax1^AA/AA^* mice were tracked by the iSCAN rodent eye tracking system while the mice were kept in dark. Relative positions of pupil centers against the position at time 0 (t_0_) were plotted in the oscillograms. **(B)** Peak positions that the pupil centers moved at vertical and horizontal axes are measured and the averages are shown in graphs. Error bars are SD (n=6). **(C)** The P45 mice were adapted in dark for 30 mins and illuminated with room light. **(E)** Alternatively, the dark-adapted mice were illuminated monocularly with a point light. Pupil diameters were measured by iSCAN eye tracking system before and after the illuminations. Relative pupil diameters against t_0_ are plotted in oscillograms. **(D** and **F)** Average pupil diameters of the mice in dark (t_0_ ∼ t_5_) and light (t_5_ ∼ t_14_) conditions are measured and shown in graphs. Error bars are SD (n=6 [*Vax1^+/+^*] and n=7 [*Vax1^AA/AA^*]). *, p<0.05; **,p<0.01; ***, p<0.005.

Spontaneous oscillations of *Vax1^AA/AA^* mouse eyes were also evident during pupil contraction responses to binocular light illumination (Video 19). However, the speed of pupil contraction was not significantly different between *Vax1^+/+^* and *Vax1^AA/AA^* mice (Figure 8C and 8D; Video 18 and 19). Interestingly, pupil contraction occurred faster in the consensual (i.e., unstimulated) eyes than that in direct (i.e., stimulated) eyes of *Vax1^AA/AA^* mice given monocular illumination (Figure 8C; Video 21). This was contrast to the patterns in *Vax1^+/+^* mice, which exhibited immediate contraction of pupils in direct eyes, followed by contraction of the pupils of consensual eyes (Figure 8E and 8F; Video 20). The results suggest that pupillary oculomotor outputs are sent in opposite routes in *Vax1^AA/AA^* mice compared with *Vax1^+/+^* mice.

### Vax1^AA/AA^ mice exhibit visuomotor anomalies

We also investigated the optokinetic reflex (OKR) of mouse eyes to moving objects (Cahill and Nathans, 2008). *Vax1^+/+^* mice rotate their eyes correspondingly and periodically in the direction of movement of black and white vertical stripes, which rotate clockwise or counter clockwise (Figure 9A – 9C; Video 22 and 24); however, they did not rotate their eyes when the stripes converged, diverged, or stopped in the front eye field (Figure 9A – 9C; Video 26, 28, and 30). Interestingly, the eyes of *Vax1^AA/AA^* mice stopped moving when the stripes moved clockwise or counterclockwise (Figure 9A – 9C; Video 23 and 25), whereas they exhibited see-saw nystagmus in response to other stimuli (Figure 9A – 9C; Video 27, 29, and 31). These results suggest that *Vax1^AA/AA^* mice might recognize the movement of objects; however, their visuomotor systems do not operate in the same way as they do in *Vax1^+/+^* mice, which rotate their eyes and heads correspondingly to moving objects.

**Figure 9.**
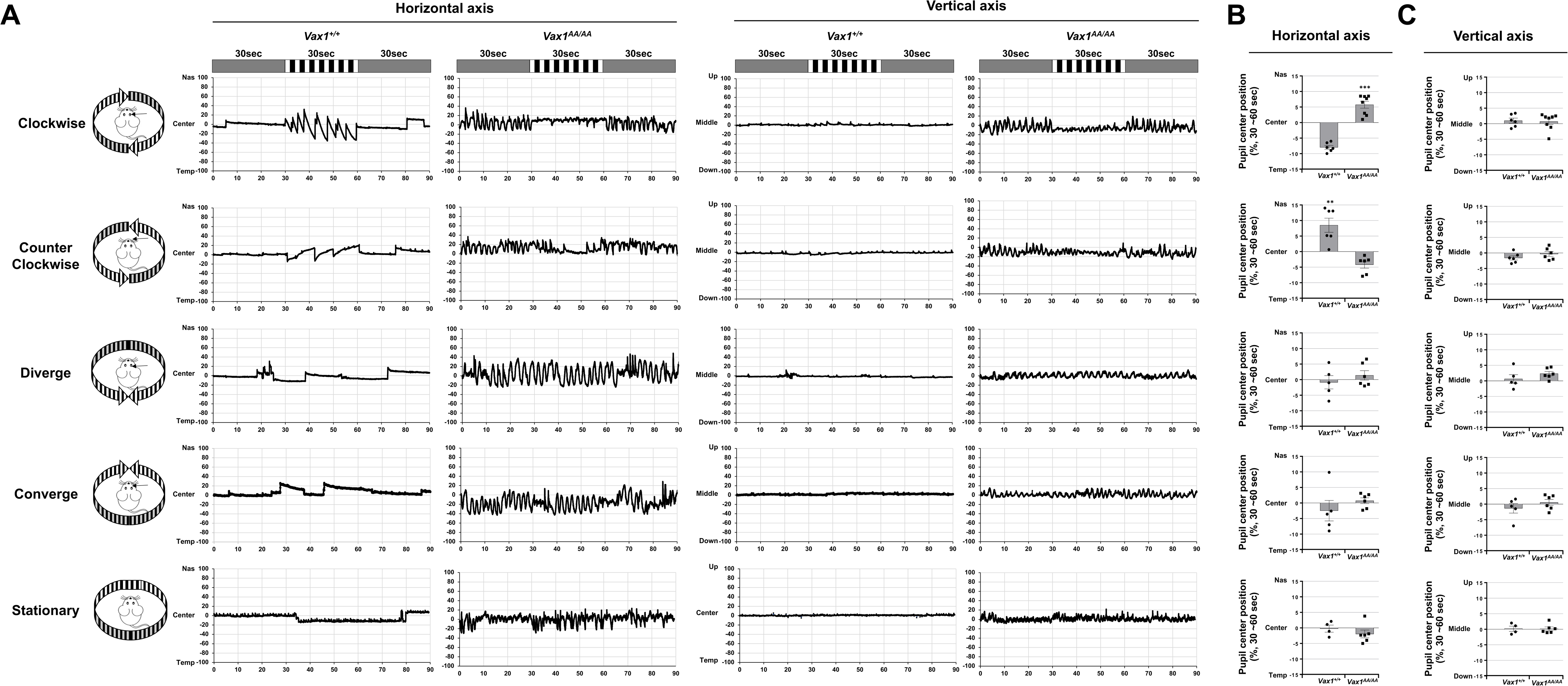
Visuomotor anmalies of *Vax1^AA/AA^* mice. **(A)** Head-fixed P45 *Vax1^+/+^* and *Vax1^AA/AA^* mice were positioned in a chamber surrounded by monitors, which display gray background. Center positions of the pupils in right and left eyes were marked and then tracked by the iSCAN rodent eye tracking system while the mice were exposed to the monitors display black and white vertical stripes (0.2 c/d), which are moving in the indicated directions for 30 seconds. Relative positions of pupil centers of right eyes against the position at t0 are plotted in the oscillograms. Peak positions of the pupil centers between 30 sec and 60 sec at horizontal **(B)** and vertical **(C)** axes were collected, and the averages are shown in graphs. Error bars are SD (n=6 [*Vax1^+/+^*] and n=8 [*Vax1^AA/AA^*]).

## Discussion

Many vertebrate organs exhibit bilateral symmetry. However, these paired organs are not simply duplications, but instead are frequently functional complements of each other. For instance, the left hemisphere of the human cerebral cortex houses the language center, whereas the right hemisphere is where pattern recognition occurs (Wolman, 2012). This is demonstrated by the ‘split-brain’ phenomenon, in which an individual whose two cerebral hemispheres are not connected by the CC cannot match words to the corresponding objects. In vertebrate binocular visual systems, bilateral projection of RGC axons at the OC is necessary for the brain to receive visual information coming from the two eyes (Herrera et al., 2019; Petros et al., 2008). Therefore, proper development of the OC is required for overlapping focal-drifted images from each eye and coordinating bilateral oculomotor responses.

Agenesis of the OC (AOC) has been reported in various vertebrates, including human, dog, and fish (Apkarian et al., 1995; Dell’Osso and Williams, 1995; Karlstrom et al., 1996; Williams et al., 1994). However, the molecular features of AOC have not been identified except in *belladonna* (*bel*) zebrafishes, which carry mutations in the LIM homeobox 2 (*lhx2*) gene (Seth et al., 2006). In the achiasmatic *bel* mutants, ventral diencephalic regions, including the preoptic area (POA), vHT and OS, are not patterned properly by virtue of the failure to express key genes, including *vax2*, *zic2.1*, and *pax2.1* (Seth et al., 2006). However, Lhx2 and Pax2 are properly expressed in the *Vax1^AA/AA^* mouse OS (Figure 3C; Lhx2 data not shown). Vax2 and Zic2 are not present in the OS of *Vax1^+/+^* and *Vax1^AA/AA^* mice, but are expressed in the ventral and ventral-temporal retina, respectively (Figure 5C; Vax2 data not shown). Moreover, the OS was specified properly in *Vax1^AA/AA^* mice, whereas the OS specification in *Vax1^-/-^* mice was incomplete (Figure 3C and 3D).

However, the OS in *Vax1^AA/AA^* mice exhibited developmental delay. In these mice, the optic fissures at the OS were closed completely by E16.5, whereas in *Vax1^+/+^* mice, they had already disappeared and OS cells had spread among RGC axons in a salt-and-pepper pattern by E14.5 (Figure 3C and 3D). The morphology of the E14.5 *Vax1^AA/AA^* mouse OS was largely similar to that of E12.5 *Vax1^+/+^* mice, whose OS cells exhibited neuroepithelial characteristics (Figure 3B). The molecular mechanisms underlying the maturation of OS cells still remain largely unknown, although several markers of OS cell lineage have been identified. For instance, Nestin and E-cad are expressed in the OS neuroepithelium; S100ß is expressed in OS APCs; and glial fibrillary acidic protein (Gfap) is expressed in OS astrocytes (Tao and Zhang, 2014). These markers, however, are not mutually exclusive. S100ß can be detected together with E-cad in OS cells (Figure 3D), suggesting an OS cell transition state between neuroepithelium and APC. As a result, it is difficult to dissect developmental stages of OS cells clearly with the limited information available. Thus, a more comprehensive understanding of OS maturation will require the identification of additional markers that are selectively expressed at specific OS developmental stages.

The patterning of the HT was also unlikely affected in *Vax1^AA/AA^* mice (Figure 5C), as it had also been reported previously in *Vax1^-/-^* mice (Bertuzzi et al., 1999; Hallonet et al., 1999). However, the HT of E14.5 *Vax1^AA/AA^* mouse embryos did not exhibit normal structure seen in *Vax1^+/+^* littermates. It protrudes ventrally, whereas the vHT of *Vax1^+/+^* mice was flattened over the RGC axons (Figure 5A and 5B). The results suggest that the structural alteration of HT of *Vax1^AA/AA^* mice could be resulted from the absence of RGC axons, which might provide a platform for vHT flattening. Alternatively, the OS, which formed a continuous neuroepithelial structure with the HT and failed to close optic fissures (Figure 3B), might be also related with the vHT phenotype of E14.5 *Vax1^AA/AA^* mice. Finally, in addition to these extrinsic factors, the structural alteration could be induced by intrinsic gene expression changes in the vHT cells. However, the vHT patterning factor, *Shh*, was detected in the similar domain of *Vax1^+/+^* and *Vax1^AA/AA^* mouse HT (Figure 5C). Glast-positive radial glial cells were also detected in the vHT of *Vax1^AA/AA^* mice (Figure 5C) and expressed Vegfa and Nr-Cam (Figure 5D), which are known to support RGC axon growth toward the midline (Erskine et al., 2011; Williams et al., 2006). Therefore, the intrinsic factors, which were changed in the vHT of *Vax1^AA/AA^* mouse embryos to cause vHT malformation, need to be identified by comprehensive analyses of the vHT cells in future studies.

It has been proposed that Vax1 specifies the OS in vertebrates by directly suppressing the expression of a retinal fate determinant Pax6 (Bertuzzi et al., 1999; Mui et al., 2005). Expression of Pax6 in the OS was suppressed properly in *Vax1^AA/AA^* mice, whereas it was induced ectopically in the *Vax1^-/-^* mouse OS (Figure 3C). The ability of Vax1^AA^ to bind DNA sequences of *Pax6 α*-enhancer was not different from that of Vax1 (Figure 2E and 2F), suggesting that Vax1^AA^ could regulate target gene expression as efficiently as Vax1. These results suggest that Vax1 transcription factor activity is not crucial for OS maturation, whereas it is necessary for the specification of the OS. These also suggest that secreted Vax1 supports the OS maturation. The secreted Vax1 might function in autocrine and paracrine manners (Figure 10B). The secreted Vax1 might reenter the OS cells to induce target genes at post-transcription levels, as it did in neighboring RGC axons to induce local mRNA translation of axon growth stimulating genes (Kim et al., 2014). In addition, the OS-derived Vax1 in RGC axons might also induce the expression of the factors that trigger the maturation of OS cells. Therefore, Vax1 could play a role as a signaling factor that couples the OS maturation and RGC axon growth. Previously, RGC-derived Shh was shown to promote OS cell proliferation and differentiation (Burne and Raff, 1997; Dakubo et al., 2003), suggesting a possibility that the OS-derived Vax1 induces *Shh* mRNA translation in the axons. However, the deletion of *Shh* in the retina did not delay the closure of OS fissures (Dakubo et al., 2003). Therefore, the delayed maturation of OS cells in *Vax1^AA/AA^* mice might not be resulted from Shh reduction in neighboring RGC axons.

**Figure 10.**
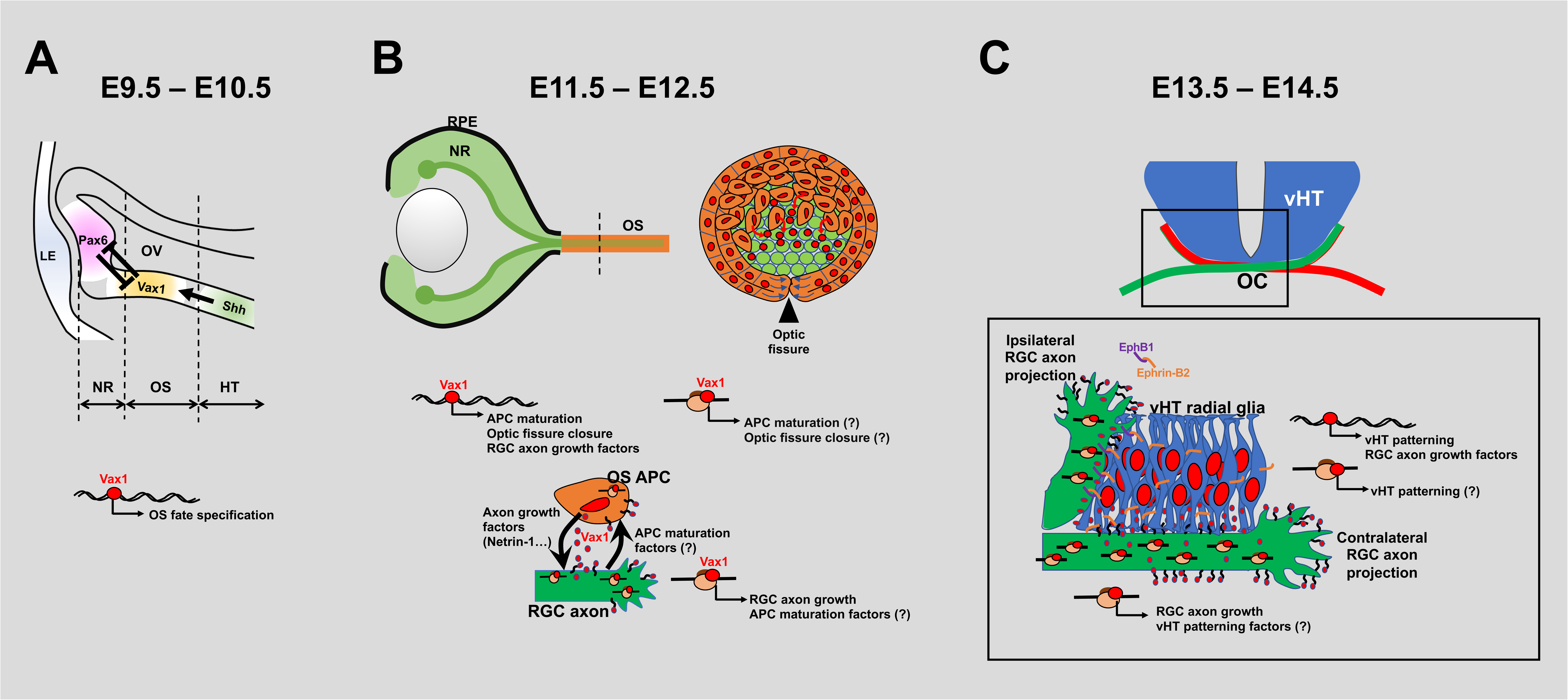
Schematic diagram depicts the pleiotropic roles of Vax1 in mouse optic nerve development. **(A)** Vax1 is expressed in the proximal optic vesicle (OV) at downstream of Shh signal, which comes from the ventral midline, specifies the OS fate by suppressing the expression of retinal determinants, including Pax6 (Mui et al., 2005), between E9.5 and E10.5. **(B)** This event is followed by the invagination of the OV that forms a double-layer optic cup of neural retina (NR) and RPE in the distal OV between E11.5 and E12.5. The fissures in the OS (arrowhead) and the optic cup are also closed as a result of the invagination. The optic fissure closure is impaired in *Vax^-/-^* mice but delayed in *Vax1^AA/AA^* mice, which express the secretion-defective and transcription-competent Vax1^AA^ mutant (Figure 3B). These suggest that Vax1 regulates the fissure closure not only through the expression of target genes in the OS cell nucleus but also via the secretion. The secreted Vax1 not only penetrates neighboring RGC axons, it can also reenter in the OS APCs. The autocrine Vax1, in turn, might induce the expression of the genes, which regulate APC maturation and/or the fissure closure, as the OS APC-derived Vax1 in RGC axons promoted the axonal growth via local mRNA translation (Kim et al., 2014). APC maturation factors could be also included in the targets of Vax1 in RGC axons. Thus, APC maturation could be also delayed in *Vax1^AA/AA^* mice, in which RGC axon growth is retarded. N, nucleus. **(C)** Vax1 could also act in the neighboring RGC axons as well as the vHT cells, as it did work in the OS. Thus, defective secretion of Vax1 could not only affect the growth of RGC axons approaching the vHT, but it could also cause developmental disorders in the vHT. Therefore, bilateral projection of RGC axons is not only dependent on the expression of pathway selection cues, including Vegfa, Nr-Cam, and Ephrin-B2, in the vHT, but it also requires proper expression of vHT patterning genes.

Collectively, our study suggests pleiotropic roles of Vax1 in mouse optic nerve development. Vax1 is expressed in mouse ventral forebrain in after E8.5 and specifies the HT and OS in the ventral diencephalic area (Bertuzzi et al., 1999; Hallonet et al., 1999). This might be mediated by transcriptional regulation of various target genes, including a negative target *Pax6* (Mui et al., 2005) (Figure 10A). In the OS, Vax1 not only induces the development of the OS cell lineage but also regulates the growth of neighboring RGC axons. The latter is not only promoted by Vax1-dependent expression of RGC axon growth factors, such as *Netrin-1* and *Sema5a* (Figure 4 – figure supplement 2), in the OS APCs, but it also requires intercellular transfer of Vax1 protein that enhances local mRNA translation in RGC axons (Kim et al., 2014) (Figure 10B). Therefore, not only the loss of Vax1 (i.e., knock-out) but also the inactivation of Vax1 transfer (i.e., KA-to-AA mutation) could result in the retarded RGC axon growth (Figure 4B and 4C). These mutations also lead to the malformation of vHT structure and defective RGC axon growth to the midline (Figure 5A and 10C). Thus, the identification of transcription and translation targets of Vax1 in the OS, vHT, and RGC axons should be granted for future studies to understand the pleiotropic roles of Vax1 in mouse optic nerve development.

One of the most prominent characteristics of achiasmatic animals is seesaw nystagmus (Apkarian et al., 1995; Dell’Osso and Williams, 1995; Williams et al., 1994). In mammalian oculomotor systems, the rostral interstitial nucleus of medial longitudinal fasciculus (riMLF) and the interstitial nucleus of Cajal (INC) in the tegmentum of the midbrain were identified as a key regulatory center for vertical gaze (Büttner et al., 2002; Zee, 1986). The riMLF neurons project directly to the oculomotor nucleus (ON) in the midbrain, and indirectly via the INC, to trigger ocular muscle contraction (Büttner et al., 2002). In *Vax1^+/+^* mice, RGC axons bilaterally innervate brain areas (Figure 6C – 6E), which send the signals to riMLF neurons. Therefore, riMLF neurons at both sides can be activated commonly even by monocular stimulation, consequently the four ocular muscles—IR, SO, IO and SR—can contract simultaneously to result in vertical gaze. In contrast, the oculomotor centers in *Vax1^AA/AA^* mice receive the visual inputs only from ipsilateral eyes (Figure 6C). Therefore, signals from right eye in *Vax1^AA/AA^* mice might induce contractions of IR and IO in the right eye and SR and SO in the left eye, resulting in nasal-downward pendular motion of the right eye and temporal-upward pendular movement of the left eye (Figure 8A; Figure 8 - figure supplement 1; Video 17). This vertically out of phase but horizontally in-phase eye movement is then followed by temporal-upward and nasal-downward pendular motions, completing one oscillation cycle. This alternating eye movement, therefore, suggests that the signals from the two eyes are desynchronized and/or feeding back on each other.

The paramedian pontine reticular formation (PPRF), a horizontal gaze center in the pons that receives oculomotor inputs from ipsilateral prefrontal cortex (Buttner-Ennever and Buttner, 1988; Zee, 1986). The PPRF also receives inputs from the contralateral SC and delivers them to the neighboring abducens nucleus (ABN). Abducens internuclear neurons (AIN) in the ABN then relay the signals to CN-III across the midline to stimulate the medial rectus (MR) ocular muscle, while abducens motor neurons (AMN) in the ABN connect ipsilaterally to the lateral rectus (LR). Given the binocular nature of efferent nerve fibers of ABN neurons, contractions of the MR in one eye and the LR in the other eye rotate both eyes in the same direction (Figure 8C). Horizontal gaze is therefore achieved when MR and LR in the same eye are activated simultaneously. However, because of the seesaw nystagmus, horizontal eye gaze was not observed in *Vax1^AA/AA^* mice.

The eyes of *Vax1^AA/AA^* mice frequently stopped oscillating when mice were presented objects moving clockwise or counterclockwise, instead of rotating correspondingly in the direction of object movement as *Vax1^+/+^* mice (Figure 9A; Video 23 and 25). This stimulus-driven ectopic gaze of *Vax1^AA/AA^* mouse eyes might result from the combinatorial activation of extraocular muscles. As it is noted above, the seesaw nystagmus of *Vax1^AA/AA^* mice is likely driven by alternating activation IR+IO and SR+SO ocular muscles. Therefore, IR+IO-driven nasal (and downward) movement of the right eye could be antagonized by LR-induced temporal movement in response to clockwise movement of the object. At the same time, SR+SO-driven temporal (and upward) movement of the left eye might be antagonized by MR-induced nasal movement. However, given the presence of the respective downward and upward forces after antagonism by the LR and MR, the right eye position is slightly below the center and the left eye position is slightly above the center during the ectopic gaze in responding to clockwise stripe rotation (Figure 9A).

The *bel rev* achiasmatic zebrafish model exhibits nystagmus when facing non-moving stripes (Huang et al., 2006). The nystagmus of the fish, however, disappears in the dark and after non-patterned illumination of the visual field, which triggers seesaw nystagmus in achiasmatic mammals (Apkarian et al., 1995; Dell’Osso et al., 1998) (Figure 8A). Furthermore, in contrast to the stimulus-driven ectopic gaze in achiasmatic *Vax1^AA/AA^* mice (Figure 9A), *bel rev* zebrafish exhibit reversed horizontal OKR in response to the rotation of stripes (Neuhauss et al., 1999; Rick et al., 2000). It was proposed that the ipsilateral RGC projection supplies a reversed retinal slip velocity input to the optokinetic system in zebrafish to elicit eye movements that compensate for retinal slip in the wrong direction (Neuhauss et al., 1999; Rick et al., 2000). However, reversed OKR and postural abnormalities have not been reported in humans (Apkarian et al., 1995), dogs (Dell’Osso and Williams, 1995; Hogan and Williams, 1995), or mice (Figure 9A). These results suggest that the oculomotor circuits of zebrafish are likely different from those in mammals.

The oculomotor circuit also controls bilateral pupillary contraction, which is triggered by RGCs that are wired to the pretectal nucleus (PN) in the midbrain (Szabadi, 2018). PN neurons relay these signals to the Edinger-Westphal nucleus (EWN), which projects axons to the ipsilateral CN-III to induce pupillary contraction (Hultborn et al., 1978; Kourouyan and Horton, 1997). It has been suggested that the EWN receives signals from ipsilateral and contralateral PNs to induce bilateral pupillary contraction. However, given the faster pupil contraction of directly stimulated eyes compared with consensual eyes in *Vax1^+/+^* mice (Figure 8E), the PN might primarily stimulate the contralateral EWN in mice. Consequently, the repeated midline crossings at retina-PN and PN-EWN axes enable the directly stimulated eye to respond faster than the consensual eye. However, an ipsilateral retina-PN connection followed by a contralateral PN-EWN connection might result in an inverse order of pupillary contraction in *Vax1^AA/AA^* mice (Figure 8E). These results suggest that the operation of the oculomotor system depends on a constant number of midline crossings; therefore, the system cannot function properly if one of those commissures is missing.

The axons of SC neurons also project contralaterally to the cervical spinal cord through the tectospinal tract, and can trigger head turns in response to visual stimuli (Gandhi and Katnani, 2011). Therefore, activation of the right SC, which receives a majority of its inputs from the left eye, which captures objects in the left visual field, predominantly contracts the left neck muscle to trigger a leftward head turn in *Vax1^+/+^* mice. Given the exclusive ipsilateral retinocollicular connection, the spinal outputs in achiasmatic *Vax1^AA/AA^* mice are likely inverse to those in *Vax1^+/+^* mice. This might make *Vax1^AA/AA^* mice turn their heads in the direction opposite the stimulus. *Vax1^AA/AA^* mice, however, startled instead of turning their heads in the direction opposite the movement of horizontally drifting stripes (Figure 7I; Video 14). These results suggest that head-turn behavior is not only determined by eye-SC-spinal cord circuits, but, given the agenesis of CC in *Vax1^AA/AA^* mice (Figure 6A), is also affected by cortical circuits that regulate activity of the PPRF to pursue the objects (Gandhi and Katnani, 2011).

Our results show that retinogeniculate and retinocollicular connections are likely remained intact in *Vax1^AA/AA^* mice (Figure 6C and 6D), although those are wired exclusively in the ipsilateral paths. Furthermore, it might be possible that their visual perceptions are also preserved by reorganizing intracortical connections as it was proposed in the achiasmatic human cases (Hoffmann et al., 2012; Sinha and Meng, 2012). These suggest the impaired OKR of *Vax1^AA/AA^* mice might be resulted from the defects in visuomotor responses triggered by ipsilaterally-biased visual inputs. However, given the absence of CC (Figure 6A) and reduced retinal activity (Figure 7A), the anomalies can be also influenced by the splitted cerebral cortex and the retina with reduced cone photoreceptor activity.

## Methods

### Mouse strains

*Vax1^-/-^* and *Pax6 α-Cre* mice were reported previously (Bertuzzi et al., 1999; Marquardt et al., 2001). *Gt(ROSA)26Sor^tm4(ACTB-tdTomato,-EGFP)Luo^/J* (*R26^tm4^*) and *Gt(ROSA)26Sor^tm11(CAG- tdTomato*,-GFP*)Nat^/J* (*R26^tm11^*) mouse strains were purchased from Jackson laboratory (Muzumdar et al., 2007; Wang et al., 2019).

*Vax1^AA/AA^* mice were generated using CRISPR/Cas9 system with a CRISPR RNA (crRNA) and a trans-activating crRNA (tracrRNA) in following procedures. Two crRNAs were designed to target two nearby sites of the second exon of *Vax1* gene, which include DNA sequences encoding K101 and R102 (Figure 1 - figure supplement 1A). The sequences of crRNAs were: #1, 5-TGGATCTGGACCGGCCCAAG-3 and #2, 5- AAGGACGTGCGAGTCCTCTT-3. The synthetic single-stranded DNA oligonucleotide (ssODN) containing missense (KR-to-AA) and synonymous mutations, 5’- TCTCAGAGAGATTGAGCTGCCGAGCCAGCTCGGTTCTCTCCCGGCCCACCACGTATTGGCAACGCTGGAACTCCATCTCCAGCCTGTAGAGCTGCTCAGCTGTAAACGAGGTCCTGGTCGCCGCGGGCCGGTCCAGATCCAAGCCTTTGGGCAGGATGATTTCTCGGATAGACCCCTTGGCATCTAGGAAAGGG-3’ (IDT, Inc., USA), was used as a donor DNA. We then injected those crRNA/tracrRNA duplex and ssODN together with Cas9 mRNA (Toolgen, Inc., Seoul, Korea) into the cytoplasm of one cell-stage *C57BL/6J* mouse embryos. To screen the founders carrying the KR-to-AA mutation in the *Vax1* gene, PCR was performed (Figure 1 - figure supplement 1B). The primer sequences of crRNAs, donor DNA, and genotyping primers are proved in Table EV1. Then, the genomic regions spanning the mutated second exon of the founder mice were validated by direct-sequencing analysis (Bionics Co., Ltd., Seoul, Korea). We obtained three *Vax1^+/AA^* male mice after analyzing 83 pubs obtained from the 439 injected embryos. The off-springs of the *Vax1^+/AA^* mouse were then backcrossed with wild-type *C57BL/6J* mice over 6 generations to eliminate unwanted off-target mutations introduced by CRISPR/Cas9. All experiments were performed according to the Korean Ministry of Food and Drug Safety (MFDS) guidelines for animal research. The protocols were certified by the Institutional Animal Care and Use Committee (IACUC) of KAIST (KA2010-17) and Yonsei University (A-201507-390-01). All mice used in this study were maintained in a specific pathogen-free facility of KAIST Laboratory Animal Resource Center and Yonsei Laboratory Animal Research Center.

### Cell and explants culture

Human cervical cancer HeLa cells and human embryonic kidney (HEK) 293T cells were cultured in Dulbecco’s Eagle Modified Media (DMEM) supplemented with 10% fetal bovine serum (FBS). For the luciferase assay, HEK293T cells (10^5^) were transfected with 1 μg pCAGIG-V5 vectors encode Vax1, Vax1^AA^, or Vax1(R152S) cDNA together with pGL3- Tcf7l2-luciferase (0.2 μg) and pCMV-β-gal (0.2 μg) reporter constructs. Luciferase activities in the transfected cells were measured at 24h post-transfection, and normalized by β-galactosidase activities to obtain relative luciferase activity of the cells. To test cellular penetration of Vax1 protein, V5 peptides or V5-Vax1 proteins, which were purified from HEK293T cells, were added into the growth media (1.5 pmol/ml [final concentration]) of HeLa cells, which express GFP-Sdc2. Distribution of V5 peptides and V5-Vax1 proteins on cell surface and inside the cells were examined by immunostaining with mouse anti-V5 and chicken anti-GFP antibodies.

Retinal explants prepared as described previously (Kim et al., 2014). Briefly, the retina was prepared from mouse embryos at E13, and mixed with collagen in DMEM with 10% FBS. The retinal explants in collagen were then cultured in Neurobasal medium containing B27 supplement (Invitrogen Inc.) for 48h to allow the axons grow from the explants. 6X-Histidine peptides or Vax1-6X-His proteins, which were purified from E.coli, were then added into the culture medium (2 pmol/ml [final concentration]) of retinal explants. Alternatively, the retinal explants were placed next to collagen droplets containing HEK293 cells (10^5^ cells/droplet), which express Vax1 or Vax1^AA^. The lengths of retinal axons grown from the explants were measured before and after the treatments to determine axon growth rate.

Slab embryo culture with collagen gel was also performed as described previously (Kim et al., 2014). Collagen droplets mixed with Flag peptides (10 μg/ml) or Flag-Vax1 proteins (200 μg/ml) were prepared and placed into the third ventricle of the slab embryos. The embryos were then incubated for 12 h at 37°C in a humidified atmosphere supplemented with 7% CO_2_.

### Detection of Vax1 proteins in the growth medium

Heparin (10 mg/ml) was added into growth medium of HEK293T cells (10^7^) express V5- Vax1 or V5-Vax1^AA^. Macromolecules including the proteins and lipids in the growth medium were precipitated by adding trichloroacetic acid (TCA; 20% final). The precipitates were washed with cold acetone three times and dissolved in 2X-SDS sample buffer for SDS-PAGE followed by WB to detect Vax1 proteins released in the growth medium.

### Immunohistochemistry and in situ RNA hybridization

Distribution of proteins in HeLa and mouse embryonic cells were examined by immunostaining. Cultured cells were fixed in 4% paraformaldehyde (PFA) in phosphate buffered saline (PBS) for 10 mins at 36-h post-transfection. Sections of mouse embryos, eyes, and brain slabs were fixed in 4%PFA/PBS at room temperature for 2 h, and then put in a 20% sucrose/PBS solution at 4°C for 16 h before embedding in OCT (optimal cutting temperature) medium for cryofreezing and cryosection.

The cells and sections were incubated in a blocking solution containing 0.2% Triton X-100, 5% normal donkey serum, and 2% bovine serum albumen (BSA) in PBS for 1 h. To stain the proteins in the cells, the samples were incubated with the indicated primary antibodies in blocking solution without 0.2% Triton X-100 at 4°C for 16 h and then with the appropriate secondary antibodies conjugated with fluorophores. Immunofluorescence was subsequently analyzed using Olympus FV1000 and Zeiss LSM810 confocal microscopes. Antibody information is provided in Table S1.

Distributions of mRNA of interest in the embryonic sections were detected by *in situ* hybridization (ISH) with digoxygenin (DIG)-labeled RNA probes and visualized them by immunostaining with alkaline phosphatase (AP)-conjugated α-DIG followed by AP-mediated colorization, as it was described in a previous report (Kim et al., 2014).

### Tissue clearing, lightsheet microscopy, and 3D image reconstitution

To visualize tdTom* fluorescence signals, the embryos were cleared by stabilization under harsh conditions via intramolecular epoxide linkages to prevent degradation (SHIELD) method as described previously (Park et al., 2019). In brief, mouse embryos were serially incubated in SHIELD perfusion solution, SHIELD-OFF solution, and SHIELD-ON solution. The samples were then delipidated for 3-5 days at 47°C in SDS clearing buffer, followed by washing at 37°C in PBS containing 1% TritonX-100 and 0.02% sodium azide for 24h. The delipidated samples were incubated in optical clearing solution/PBS (50:50) for the incubation at RT for 12h, and then in optical clearing solution at RT for 12-24h until samples became transparent. The samples were embedded in 1.5% agarose in optical clearing solution for the imaging by Zeiss Lattice Lightsheet 7 microscope. Collected images were processed for stitching and 3D reconstruction with ZEN software (Zeiss), then analyzed by surface tool of the IMARIS 9.3 software (Bitplane) to rendering tdTom* fluorescence signals. To quantify the fluorescence signal intensity of the tdTom*-labeld RGC axons across the optic disc head and hypothalamic midline, valid fluorescence spots were identified by background subtraction. Distance between each spot and the sagittal plane was calculated.

### Chromatin immunoprecipitation (ChIP) and PCR

Chromatin immunoprecipitation was done as it was described previously (Mui et al., 2005). E10.5 mouse embryonic heads were isolated and chopped into small pieces in prior to the incubation in 1% formaldehyde in PBS at room temperature for 10 min. The nuclei were isolated for the immunoprecipitation with rabbit anti-Vax1 antibody or pre-immune rabbit IgG. DNA fragments coprecipitated with the antibodies were purified by phenol/chloroform/isoamyl alcohol extraction, and 100 ng of these immunoprecipitated DNAs were used as templates for PCR amplification of the Pax6 α-enhancer.

### Electroretinogram (ERG)

Mice were either dark- or light-adapted for 12 h before ERG recording and anesthetized with 2,2,2-tribromoethanol (Sigma). After the pupils of the mice were dilated by 0.5% tropicamide, a gold-plated objective lens was placed on the cornea and silver-embedded needle electrodes were placed at the forehead and tail. The ERG recordings were performed using Micron IV retinal imaging microscope (Phoenix Research Labs) and analyzed by Labscribe ERG software according to the manufacturer’s instruction. To obtain scotopic ERG a- and b-waves, a digital bandpass filter ranging from 0.3 to 1,000 Hz and stimulus ranging from −2.2 to 2.2 log(cd·s m^−2^) were used. To yield photopic ERG a- and b-waves, filter ranging from 2 to 200 Hz and stimulus ranging from 0.4 to 2.2 log(cd·s m^−2^) with 1.3 log(cd·s m^−2^) background were used.

### In vivo extracellular recording and data analysis

We performed *in vivo* extracellular recordings in the monocular V1 (bregma, -3.50 mm; lateral, 2.50 mm; depth, 0.70 mm) of Vax1^+/+^ and Vax1^AA/AA^ mice. Mice were anesthetized with the urethane (2 g per kg body weight, intraperitoneal injection) and restrained in a custom-designed head-fixed apparatus. A small craniotomy with the diameter of ∼0.5 mm was made over V1 of the left and the right hemispheres, and we inserted a 32-channel silicon electrode (A1x32-Poly3-10mm-50-177-CM32, Neuronexus) using micro-drive motorized manipulator (Siskiyou). After waiting 20 ∼ 30 mins for stabilization, we started recording visual responses by presenting a full-field flashing light for 5 times to the left eye. The visual stimuli were presented at 10 Hz for 500 ms, 5 pulses of 50 ms duration, and total 30 trials through a gamma-corrected monitor. Extracellular signals were filtered between 500∼5000 Hz at 30 kHz sampling rate, amplified by miniature digital head-stage (CerePlex μ, Blackrock Microsystems), and saved through data acquisition system (CerePlex Direct, Blackrock Microsystems). We performed spike sorting using the Klusters software (http://neurosuite.sourceforge.net/) and further analyzed firing rates of isolated single units using MATLAB. We analyzed the z-score of firing activity in each single unit from -1 to +2 s of the onset of the visual stimuli and plotted peri-stimulus time histogram (PSTH) of the normalized activity. Firing rate change index (FR index) of individual cells was calculated using the following formula:

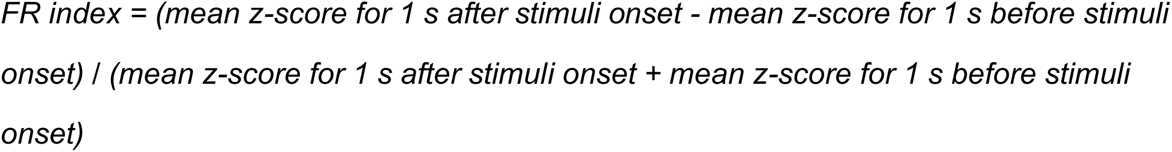

### Light-dark chamber assay

The light-dark test apparatus was composed of light (21cm(width, W) X 29cm(depth, D) X 20cm(height, H), 700 lux) and dark (21cm(W) X 13cm(D) X 20cm(H), ∼5 lux) chambers. The dark chamber is separated from light chamber by an entrance in the middle wall (5cm(W) X 8cm(H)). Mice were introduced in the light chamber with their heads toward the opposite side of the dark chamber and allowed to freely explore the apparatus for 10 min. Amounts of time spent in the light and dark chambers and number of transitions were analyzed by Ethovision XT10 software (Noldus).

### Looming assay

Looming test was performed as described previously with some modifications (Yilmaz and Meister, 2013). Briefly, the behavioral arena was prepared with an open-top acryl box (30cm(W) X 30cm(D) X 30cm(H)), which contains a nest in the shape of a triangular prism (10cm(W) X 12cm(D) X 10cm(H)). The looming disk was programmed as a black circle in a gray background, increasing its size from 2 degrees of visual angle to 20 degrees in 250 ms and maintained for 250 ms. The pattern was repeatedly presented 10 times with 500 ms of interval for each trial.

### Optomotor response (OMR)

Mouse visual acuity was measured with the OptoMotry system (Cerebral Mechanics) as previously described (Prusky et al., 2004). Mice were adapted to ambient light for 30 mins and then placed on the stimulus platform, which is surrounded by four computer monitors displaying black and white vertical stripe patterns. An event that mice stopped moving and began tracking the stripe movements with reflexive head-turn was counted as a successful visual detection. The detection thresholds were then obtained from the OptoMotry software.

### Measurements of pupillary contraction and optokinetic response (OKR)

Mouse heads were mounted to a plate and clamped to a holder to prevent head movement during measurement. Images of a mouse eye that show the pupil and the corneal reflection were recorded by CCD camera (120/240 Hz) with Infrared (IR) filter (ISCAN Inc.). To measure the OKR, the head-fixed mice were put in front of the screens that display gray background or black and white vertical stripes (30% contrast) moving at a spatial frequency of 0.2 c/d and angular velocity of 12 d/s. To examine the pupil contraction, the mice were kept in the dark for 30 secs and then exposed to 500 lux of light for 10 secs. The pupil position and diameter were measured by the ISCAN software (ISCAN Inc.).

### Statistical analyses

Statistical tests were performed using Prism Software (GraphPad; v7.0) measurement tools. All data from statistical analysis are presented as the average ± STD. Comparison between two groups was done by unpaired Student’s t-test, and the differences among multiple groups were determined by analysis of variance (ANOVA) with Tukey’s post-test used to determine the significant differences among multiple groups. P-values (*p*) < 0.05 were considered as statistically significant results.

## Acknowledgements

We thank to Ilsong Choi and Anna Shin for helping mouse ocular behavior tests. This work was supported by the National Research Foundation of Korea (NRF) grants (2018R1A5A1024261) funded by Korean Ministry of Science and ICT (MSIT), South Korea. This study was performed in strict accordance with the recommendations in the Guide for the Care and Use of Laboratory Animals of the Korean Ministry of Agriculture, Food, and Rural Affairs. All of the animals were handled according to approved institutional animal care and use committee (IACUC) protocols (#13-130) of Korea Advanced Institute of Science and Technology.

## Author contributions

K.W.M. and J.W.K. wrote the manuscript; K.W.M., N.K, J.H.L., Y.S, M.K., E.J.L., J.H.K., J.L, J.M.K., J.M.Y. designed and performed the experiments and analyzed the data; J.C.L. provided an experimental model and platform; Y.G.P., S.H.L., and H.W.L., and J.W.K. supervised the project.

## Video information

Video 1. Video presentation of 3D imaging data of RGC axons in E12.5 *Vax1^+/+^* mice.

Video 2. Video presentation of 3D imaging data of RGC axons in E12.5 *Vax1^AA/AA^* mice.

Video 3. Video presentation of 3D imaging data of RGC axons in E13.5 *Vax1^+/+^* mice.

Video 4. Video presentation of 3D imaging data of RGC axons in E13.5 *Vax1^AA/AA^* mice.

Video 5. Video presentation of 3D imaging data of RGC axons in E14.5 *Vax1^+/+^* mice.

Video 6. Video presentation of 3D imaging data of RGC axons in E14.5 *Vax1^AA/AA^* mice.

Video 7. Response of *Vax1^+/+^* mice to a looming shadow.

Video 8. Response of *Vax1^AA/AA^* mice to a looming shadow.

Video 9. Response of *Pde6b^rd1/rd1^* mice to a looming shadow.

Video 10. Cliff assay of *Vax1^+/+^* mice.

Video 11. Cliff assay of *Vax1^AA/AA^* mice.

Video 12. Cliff assay of *Pde6b^rd1/rd1^* mice.

Video 13. Optomotor response of *Vax1^+/+^* mice.

Video 14. Optomotor response of *Vax1^AA/AA^* mice.

Video 15. Optomotor response of *Pde6b^rd1/rd1^* mice.

Video 16. Eye movement of *Vax1^+/+^* mice in dark.

Video 17. Eye movement of *Vax1^AA/AA^* mice in dark.

Video 18. Pupil contraction of *Vax1^+/+^* mice after binocular light illumination.

Video 19. Pupil contraction of *Vax1^AA/AA^* mice after binocular light illumination.

Video 20. Pupil contraction of *Vax1^+/+^* mice after monocular light illumination.

Video 21. Pupil contraction of *Vax1^AA/AA^* mice after monocular light illumination.

Video 22. Eye movement of *Vax1^+/+^* mice in response to vertical stripes rotating clockwise direction.

Video 23. Eye movement of *Vax1^AA/AA^* mice in response to vertical stripes rotating clockwise direction.

Video 24. Eye movement of *Vax1^+/+^* mice in response to vertical stripes rotating counter clockwise direction.

Video 25. Eye movement of *Vax1^AA/AA^* mice in response to vertical stripes rotating counter clockwise direction.

Video 26. Eye movement of *Vax1^+/+^* mice in response to converging vertical stripes.

Video 27. Eye movement of *Vax1^AA/AA^* mice in response to converging vertical stripes.

Video 28. Eye movement of *Vax1^+/+^* mice in response to diverging vertical stripes.

Video 29. Eye movement of *Vax1^AA/AA^* mice in response to diverging vertical stripes.

Video 30. Eye movement of *Vax1^+/+^* mice in response to stationary vertical stripes.

Video 31. Eye movement of *Vax1^AA/AA^* mice in response to stationary vertical stripes.

---

**Figure 2 – figure supplement 1.**
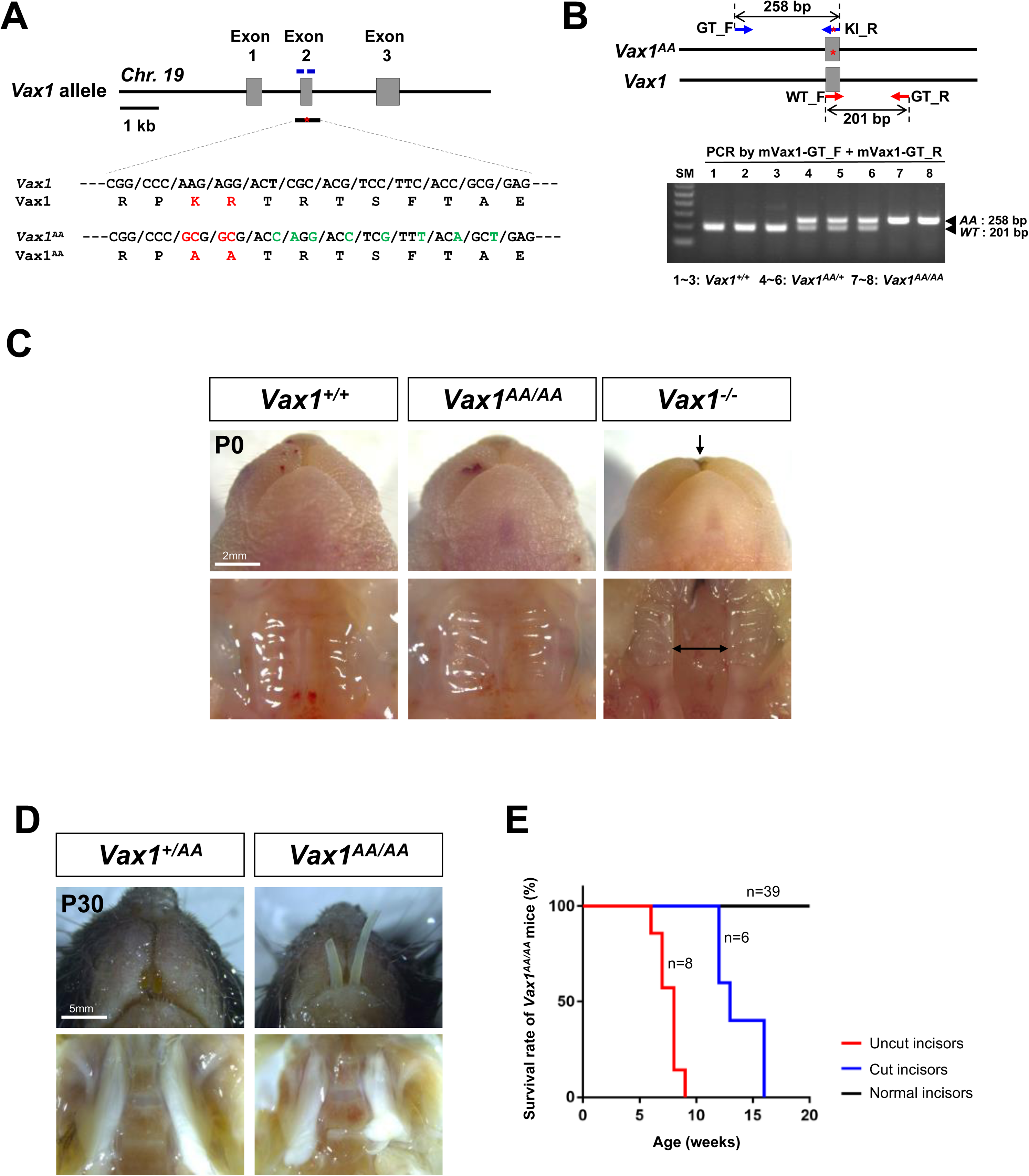
Generation of *Vax1^AA/AA^* mice. **(A)** Structure of mouse *Vax1^AA^* gene (top) and its partial sequences of exon 2 that encodes AA replacing KR in the GAG binding motif (bottom). The substituted nucleotides for missense (red, KR to AA) and synonymous (green) mutations are shown. The amino acid sequences from the Vax1 and the Vax1^AA^ alleles are shown at the bottom of each nucleotide sequence. The relative positions of two crRNAs and ssODN are indicated in short blue bars and black bar with asterisk (*), respectively. **(B)** Top, relative positions of genotyping primers are indicated by arrows. The red asterisks (*) indicate KR to AA mutation. Bottom, PCR genotyping results using genotyping primers. **(C)** Ventral views of P0 mouse mouths (top row) and palates (bottom row). Arrows indicate the clefts of lip (top row) and palate (bottom row), respectively. **(D)** Ventral views of P30 mouse mouths (top row) and palates (bottom row). **(E)** Survival rates of *Vax1^AA/AA^* mice by the indicated ages in x-axis are provided. Numbers of mice examined were provided in the graph.

**Figure 4 – figure supplement 1.**
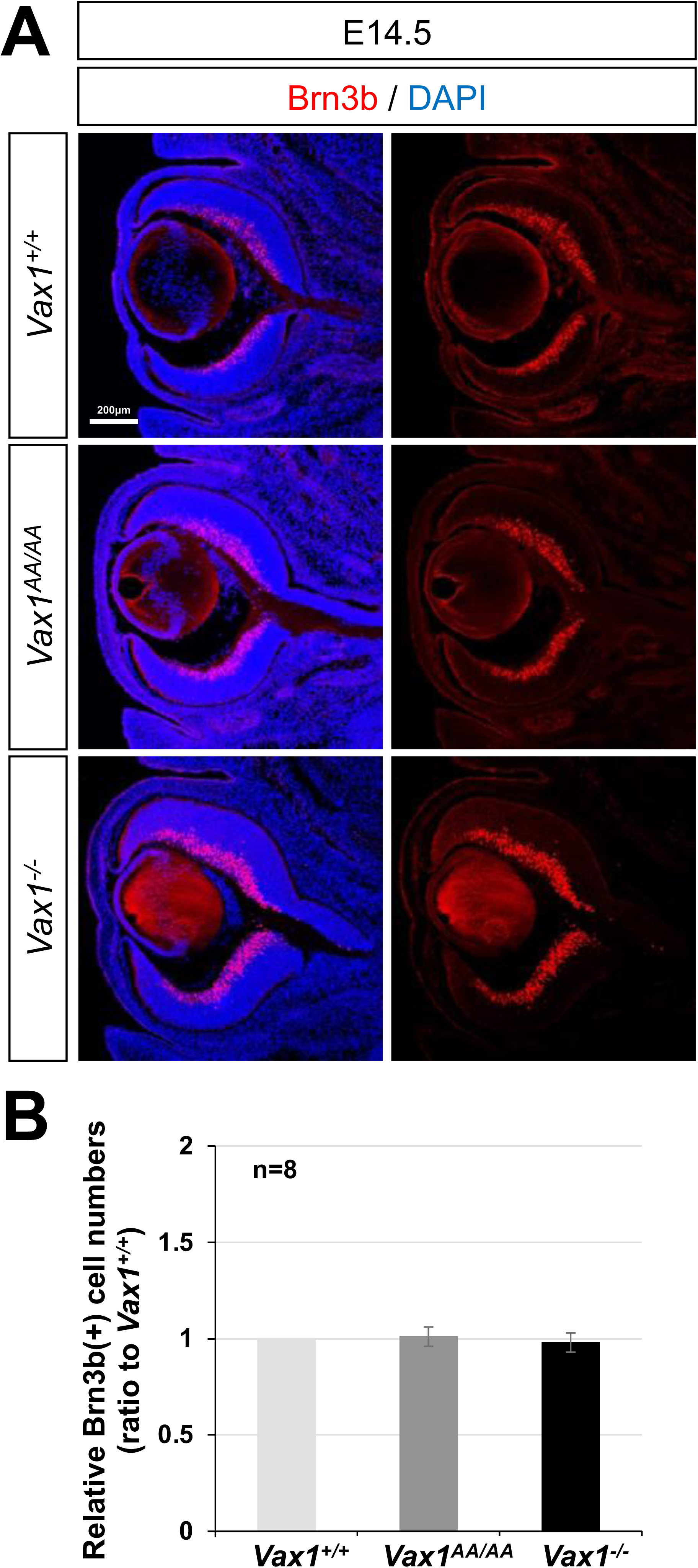
Development of Brn3b-positive RGCs in *Vax1^+/+^* and *Vax1^AA/AA^* mouse retina. (A) Distribution of RGCs in E14.5 mouse embryonic sections was determined by immunostaining of an RGC-specific marker, Brn3b. Nuclei of the cells in the sections were visualized by DAPI staining. (B) Relative numbers of Brn3b(+) RGCs in the sections are shown in the graph. The values are SD (n=8; 5 independent litters).

**Figure 4 – figure supplement 2.**
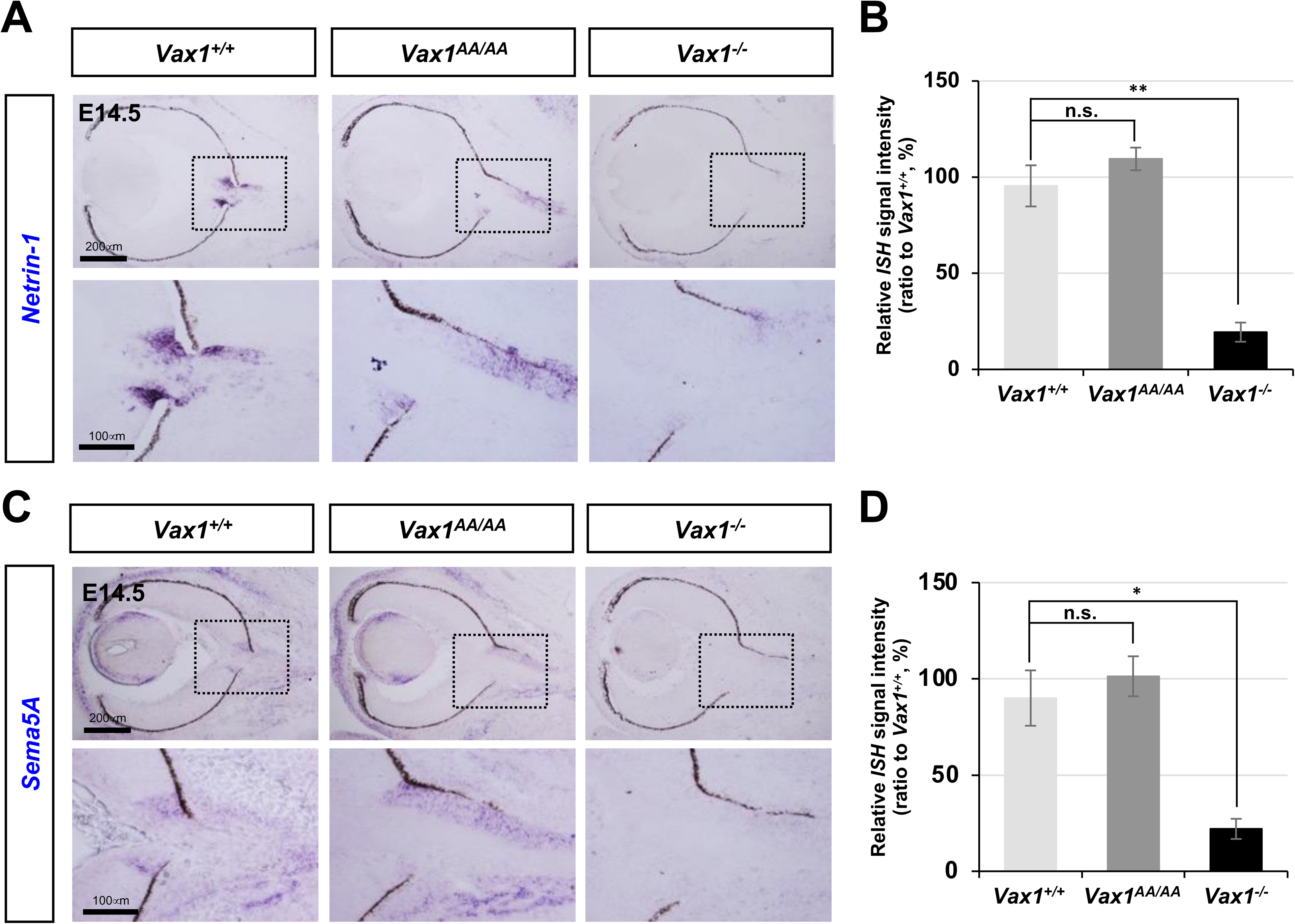
Expression of *Netrin-1* and *Sema5A* in *Vax1^+/+^* and *Vax1^AA/AA^* mouse OS. Expressions of *Netrin-1* (A) and *Sema5A* (C) mRNA in E14.5 mouse embryos with indicated genotypes were investigated by ISH. Images in the second and fourth rows are the magnified versions of boxed areas in the first and third rows. Intensities of *Netrin-1* (B) and *Sema5A* (D) ISH signals were quantified and relative values are shown in the graphs. The values are SD (n=4; 3 independent litters). *, p<0.05; **, p<0.01; n.s., not significant.

**Figure 5 – figure supplement 1.**
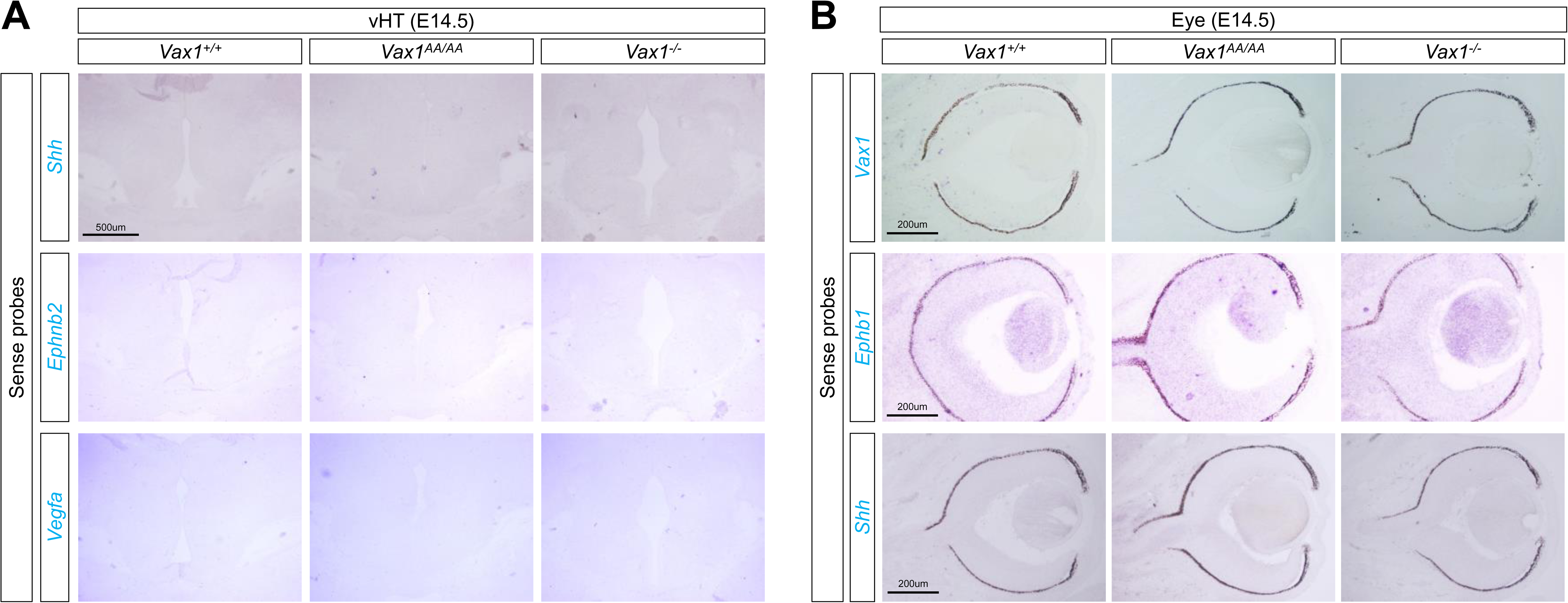
The *ISH* signals of sense probes. **(A)** Specificities of the *ISH* signals detecting *Shh* (Figure 5C), *Ephnb2* (Figure 5D), and *Vegfa* (Figure 5D) in the vHT of mouse embryonic sections were determined by the *ISH* with sense probes for the corresponding genes. **(B)** Specificities of the *ISH* signals detecting *Vax1* (Figure 2B), *Ephb1* (Figure 5E), and *Shh* (Figure 5E) in the mouse retinas were determined by the *ISH* with sense probes for the corresponding genes.

**Figure 7 – figure supplement 1.**
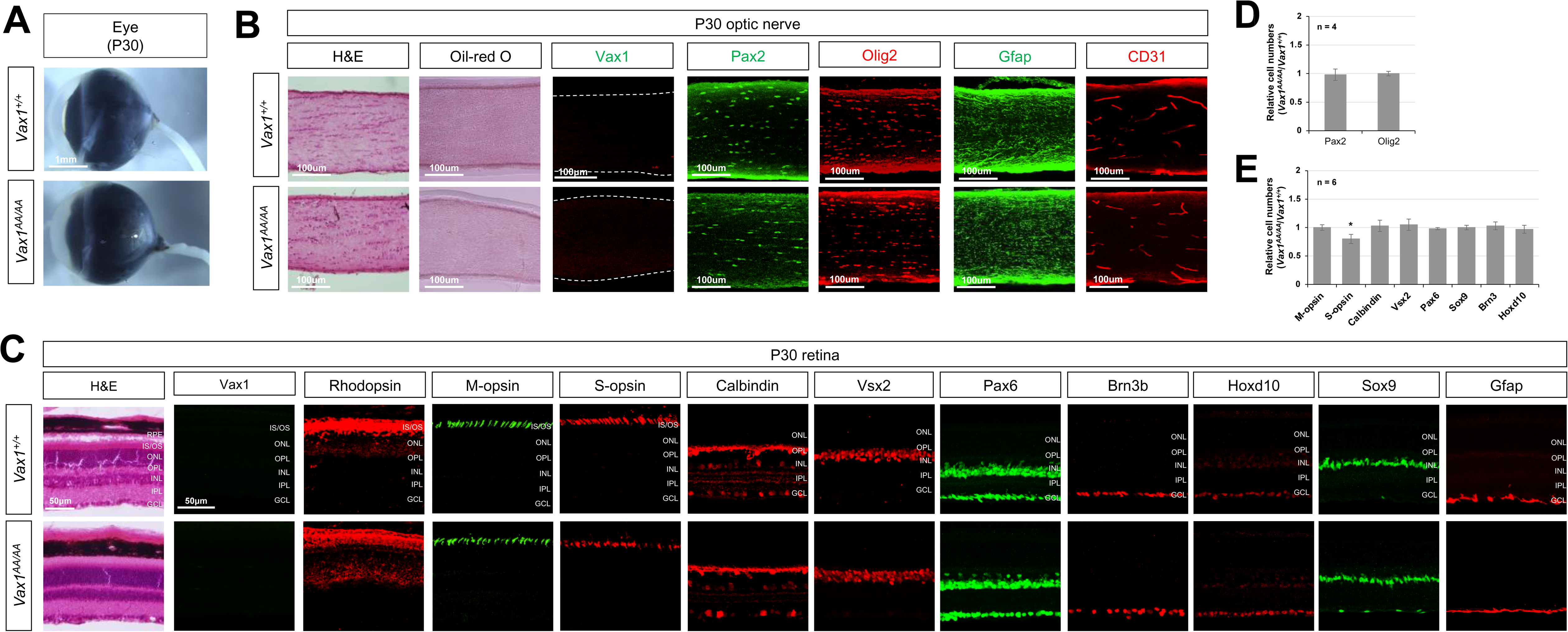
Anatomical features of the retina and optic nerve in *Vax1^+/+^* and *Vax1^AA/AA^* mice. **(A)** Lateral views of the eyes of P30 *Vax1^+/+^* and *Vax1^AA/AA^* littermate mice. **(B)** Longitudinal sections of optic nerves of P30 *Vax1^+/+^* and *Vax1^AA/AA^* littermate mice were stained by H&E and Oil Red-O to visualize the distribution of the cells and lipids in the nerves, respectively. The sections were also immunostained with the antibodies that recognize the indicated marker proteins. Pax2 and Gfap, astrocytes; CD31, endothelial cells; Olig2, oligodendrocytes. **(C)** Sections of the eyes of P30 *Vax1^+/+^* and *Vax1^AA/AA^* littermate mice were stained by H&E or the antibodies that recognize the indicated marker proteins. Rhodopsin, rod photoreceptors; M-opsin, M-cone photoreceptors; S-opsin, S-cone photoreceptors; Calbindin, horizontal cells and amacrine cell subset; Vsx2, bipolar cells; Pax6, amacrine cells; Brn3b, RGCs; Sox9, Müller glia; Gfap, astrocytes. **(D)** Relative numbers of Pax2-positive astrocytes and Olig2-positive oligodendrocytes are shown in a graph. **(E)** Relative numbers of retinal cells expressing the corresponding markers in *Vax1^AA/AA^* mice against *Vax1^+/+^* littermate mice are shown in the graph. Error bars denote SD. Numbers of samples (from independent litters) are provided in the graphs. *, *p*<0.05.

**Figure 8 – figure supplement 1.**
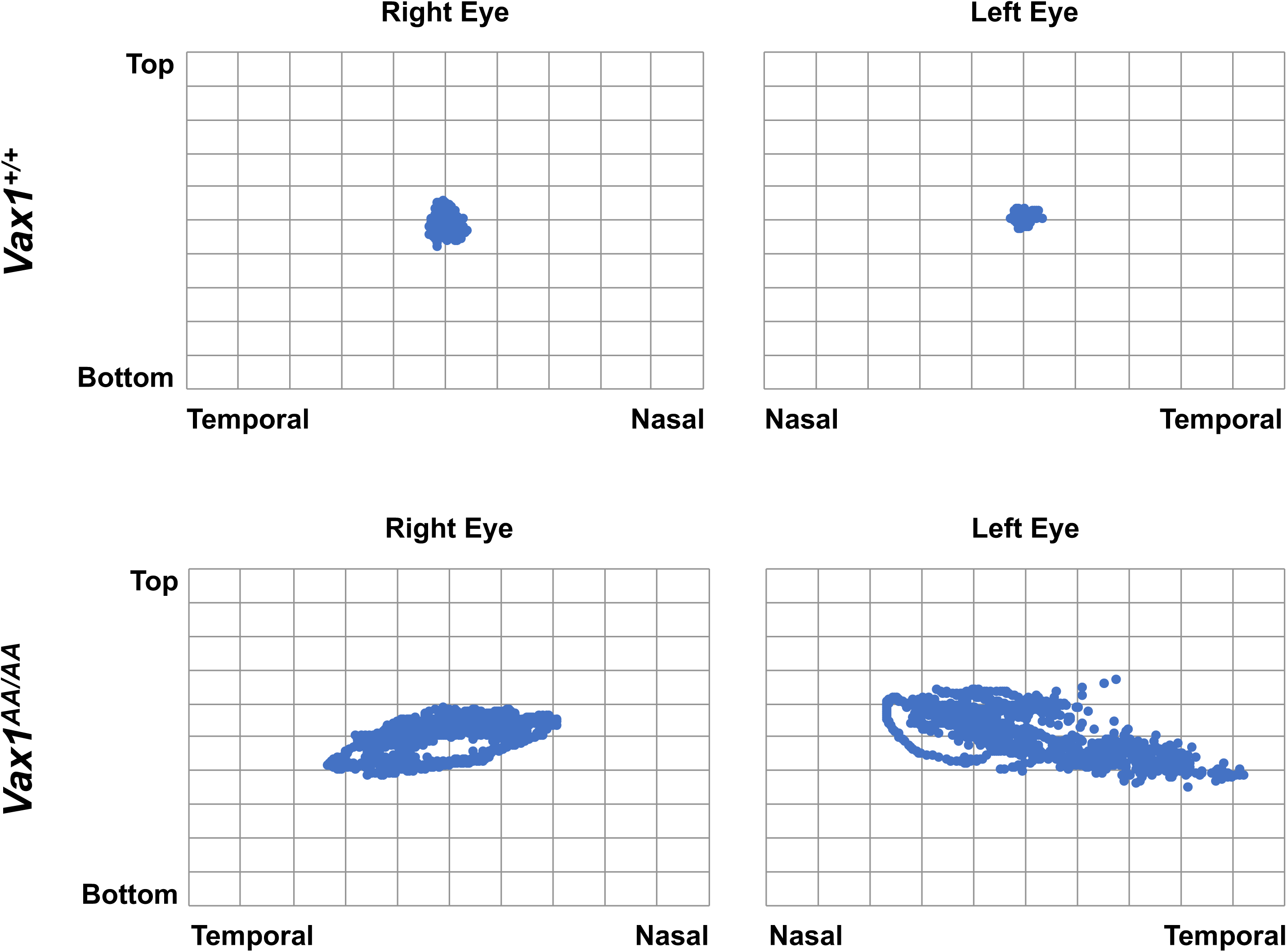
Spontaneous movement of *Vax1^AA/AA^* mouse eyes. Relative positions of the pupil centers in right and left eyes of head-fixed P45 *Vax1^+/+^* and *Vax1^AA/AA^* were recorded by the iSCAN rodent eye tracking system at every 8 msec for 10 secs while the mice were kept in dark. The positions were plotted in the graphs. The results show the eyes of *Vax1^AA/AA^* rotate spontaneously in an oval track while *Vax1^+/+^*mouse eyes keep their positions.

